# Sterile Inflammation Alters Neutrophil Kinetics in Mice

**DOI:** 10.1101/2021.02.12.430891

**Authors:** Alakesh Singh, Thiruvickraman Jothiprakasam, Jayashree V. Raghavan, Siddharth Jhunjhunwala

**Affiliations:** Centre for BioSystems Science and Engineering, Indian Institute of Science, Bengaluru, India – 560012

**Keywords:** Biomaterials, Innate immunity, Foreign-body response, Granulopoiesis, Granulocytes

## Abstract

Neutrophils play a crucial role in establishing inflammation in response to an infection or injury, but their production rates, as well as blood and tissue residence times, remain poorly characterized under these conditions. Herein, using a biomaterial implant model to establish inflammation followed by *in vivo* tracking of newly formed neutrophils, we determine neutrophil kinetics under inflammatory conditions. To obtain quantifiable information from our experimental observations, we develop an ordinary differential equation-based mathematical model to extract kinetic parameters. Our data show that in the presence of inflammation resulting in emergency granulopoiesis-like conditions, neutrophil maturation time in the bone marrow and half-life in the blood reduces by about 40%, compared to non-inflammatory conditions. Additionally, neutrophil residence time at the inflammatory site increases by two-fold. Together, these data improve our understanding of neutrophil kinetics under inflammatory conditions, which could pave the way for therapies that focus on modulating in vivo neutrophil dynamics.

## INTRODUCTION

Neutrophils are short-lived cells of the immune system that play an essential role in pathogen clearance and foreign body responses(1). They are continually produced in the bone marrow through granulopoiesis, a process where hematopoietic stem cells (HSCs) differentiate to mature neutrophils in sequential steps involving numerous intermediate progenitor populations(2, 3). Mature neutrophils are recruited into circulation where their numbers remain relatively stable throughout an individual’s lifetime(3, 4). A delicate balance between progenitor proliferation, maturation time in the bone marrow, half-life in the blood, and clearance at various tissue sites, ensure that the numbers of neutrophils in circulation remain stable(5, 6).

The kinetics of neutrophil production, maturation, circulation, and death have been extensively studied in humans and mice. In humans, over 10^11^ neutrophils are produced every day, and the maturation time in the bone marrow and half-life in circulation has been measured to be about 6 days and 11-19 hours, respectively(7–9). In mice, the number of neutrophils produced is about 10^7^ per day, their maturation time in the bone marrow is 2-3 days, half-life in circulation is 10-13 hours, and half-life at tissue sites range from 8-18 hours(6, 9–12). These measurements have been made under steady-state, non-inflammatory conditions. Under inflammatory conditions, such as during an infection, chronic disease, injury, or biomaterial implantation, neutrophil lifespans remain unknown(13).

Herein, we use a mouse model of biomaterial implantation to determine the time for maturation of neutrophils in the bone marrow, and blood and tissue residence time under sterile inflammatory conditions. We track neutrophils spatially and temporally *in vivo* and mathematically model the process to find that, in the presence of an inflammatory stimulus that results in a state resembling emergency granulopoiesis, many parameters associated with neutrophil kinetics are altered.

## MATERIALS AND METHODS

### Animal Studies

All animal studies were conducted under the Control and Supervision Rules, 1998, of the Ministry of Environment and Forest Act (Government of India), and the Institutional Animal Ethics Committee, IISc. Experiments were approved by the Committee for Purpose and Control and Supervision of Experiments on Animals (permit numbers CAF/ethics/546/2017 and CAF/ethics/718/2019). Animals were procured from the Central Animal Facility, IISc or Hylasco Biotechnology (India) Pvt. Ltd, Hyderabad, India (a Charles River Laboratory licensed supplier). All experiments were performed on 8-14-week-old (weighing 20 – 30 grams) C57BL6 mice (both female and male).

### Alginate and Chitosan microspheres

Alginate and chitosan microspheres were prepared using a Spraybase® Electrospray system in a sterile enclosure. Alginate microspheres - SLG20 alginate (Nova Matrix, FMC BioPolymer, Drammen, Norway) was dissolved in 0.86% NaCl (saline) to a concentration of 1.4 % (w/v). This solution was then passed through a 26G blunt needle, at the pressure of 0.5 bar and a voltage of 5 kV, into a 50 mM BaCl_2_ cross-linking solution. The collector distance from the tip of the needle was 5 cm. Chitosan microspheres -medium molecular weight chitosan (448877, Sigma-Aldrich, USA) was dissolved in 1 % (v/v) acetic acid solution at a concentration of 1% (w/v) and stirred overnight at room temperature (RT). The chitosan solution was then passed through a 26G blunt needle at 0.8 bar pressure and a voltage of 6 kV, into 0.9 % (w/v) sodium tripolyphosphate cross-linking solution (pH 5.0). The collector distance from the tip of the needle was 7 cm. All the microspheres were collected and washed six times with sterile saline. The microspheres were then incubated in 1% NaOH at RT, for two hours with a replacement of NaOH solution at one hour, followed by incubation in a 1.6% (v/v) paraformaldehyde solution in phosphate-buffered saline at RT for 1 hour. All microspheres were then washed with excess saline and distributed in 1.5 ml sterile microcentrifuge tubes and sealed for use in implantation procedures.

### Surgical implantation of microspheres

Microspheres were surgically implanted into the PC of mice as described(14). In brief, microspheres (450 µl) were suspended in 450 µl sterile saline for implantation. The entire volume (900 µl) was then implanted into the PC following a laparotomy procedure. In controls (also called mock controls), the animals went through the surgical procedure and were injected with 900 µl sterile saline.

### *In vivo* labelling of cells

EdU was purchased from Carbosynth (UK). EdU was dissolved in sterile saline at a concentration of 0.625 % (w/v), and each mouse received a concentration of 25 mg/Kg body weight(15) through intraperitoneal injection using a 26G needle. Surgical procedures are also an inflammatory stimulus. To exclude the inflammatory effects of surgery, we waited five days post-surgery to inject EdU. At day 5, to label proliferating cells, 5-ethynyl-2-deoxyuridine (EdU) was injected intraperitoneally (**Fig. S1**). Following EdU administration, mice were euthanized at 24-hour intervals, and labeled neutrophils were quantified using flow cytometry.

### Retrieval of cells and microspheres

Mice were anesthetized using ketamine solution to retrieve cells and microspheres. After anaesthesia, about 500-1000 μl blood was collected through the retro-orbital vein and cardiac puncture. Next, mice were euthanized by cervical dislocation. Immediately following euthanasia, 5 ml of cold phosphate-buffered saline (PBS) with 4mM EDTA (PBS-EDTA) was injected into the PC using a 26G needle. Through a small incision in the peritoneal wall, the fluid containing cells was retrieved, passed through a 100 μm filter (to filter out implants) and stored on ice before analysis. Microcapsules were collected on the 100 μm filter by rinsing the PC with PBS and stored in 1.6% paraformaldehyde for analysis. Subsequently, the spleen and a tibia and femur (to get bone marrow) were collected. A cell suspension was prepared by mincing the tissue using forceps and passing the cell solution through a 100 μm filter. Cells isolated from all sites (peritoneal fluid, blood, bone marrow, and spleen) were subjected to RBC lysis and then counted manually using a Bright-Line™ hemocytometer (0.1 mm).

### Flow cytometry

For the quantification of labeled (EdU positive) neutrophils, after counting cells from each site, 0.5 x 10^6^ cells were used for antibody staining and flow cytometry. First, cells were stained with BD Horizon™ Fixable Viability Stain 510 for 20 mins at RT. Cells were then fixed using 1.6 % paraformaldehyde for 30 mins at RT. Following one wash with excess PBS, cells were stained to estimate EdU positive cells using Sulfo-Cyanine5 azide (Lumiprobe, USA) as per the manufacturer’s instructions. Following two additional washes with PBS, cells were stained with – the monoclonal antibody against Ly6G (clone 1A8, BD Biosciences, USA) for 30 mins at 4°C, in PBS containing 1 % BSA and 4 mM EDTA (staining buffer). Finally, cells were re-suspended in staining buffer.

To monitor neutrophil activation markers in untreated mice, 0.5 x 10^6^ cells from each site were stained immediately with a combination of the following antibodies for 30 mins at 4°C: CD11b (M1/70), Ly6G (1A8), CD45 (30-F11) CD54 (3E2), CD62L (MEL-14), CD182 (V48-2310) all purchased from BD Biosciences (USA). For *ex vivo* activation –, 0.5 x 10^6^ cells from each site were first activated using 5 µg/ml Cytochalasin B (C6762, Sigma-Aldrich, USA) for 5 mins and 5 µM fMLP (F3506, Sigma-Aldrich, USA) for 30 mins at 37°C and then stained with antibodies as mentioned above. Cells were stained with propidium iodide (2 µg/ml), and PI-positive cells were excluded to identify live cells.

To identify neutrophil progenitors, RBC were lysed and 2 x 10^6^ cells from bone marrow and spleen were stained with lineage cocktail antibodies: I-A/I-E (M5/114.15.2), CD49b (HMα2), NK-1.1 (PK136), CD11c (N418), Ly6C (HK1.4), Ly6G (1A8), CD3(17A2), CD127 (A7R34), CD19(6D5), CD45R/B220 (RA3-6B2) and stem/progenitor cell markers, c-Kit (2B8), Sca-1(D7), CD34(SA376A4) and CD16/32(93). Cells were stained with propidium iodide (2 µg/ml), and PI-positive cells were excluded to identify live cells.

For all protocols, appropriate single color (using compensation beads, BD Biosciences) and fluorescence-minus-one (FMO) controls (using cells) were used to compensate the data and gate positive populations, respectively. Flow cytometry data were collected using a BD FACSCelesta (Becton Dickinson, USA) and analyzed using FlowJo (Tree Star, Ashland, OR, USA)

### Myeloperoxidase (MPO) and elastase activity assays

The following numbers of cells were used for these assays: bone marrow – 2 x 10^6^, blood - 0.2 x 10^6^, peritoneal fluid - 0.2 x 10^6^. These cells were seeded in a 96-well clear flat bottom plates. PBS was then added to make up the volume of each well to 135 µl. CTAB was dissolved in PBS at a concentration of 0.5 % (w/v), and 15 µl was added to each well to lyse the cells. The cells were then incubated at 37°C for 30 mins. After the incubation period, the plate was centrifuged at 1000 RCF at 4°C for 10 min. to pellet down all the cell debris. Supernatant aliquots (100 μl) were placed in 0.5 ml protein low bind tubes and stored at -80°C.

#### MPO activity

Supernatants were used at different dilutions for measuring MPO activity. Diluted supernatant (75 µl total volume) from each sample was added to a 96-well transparent flat bottom plate. 75 µl of 3,3′,5,5′-tetramethylbenzidine (TMB, T4444, Sigma-Aldrich, USA) was then added to each well for 90 seconds, followed by the addition of 150 µl of 1 M H_2_SO_4_ to stop the reaction. The samples were then read on a 96 well plate reader at 450 nm. Standard samples with different myeloperoxidase concentrations (from human polymorphonuclear leucocytes, 475591, Sigma-Aldrich, USA) were prepared to obtain a standard curve in the range of 26-1333 mU/ml. MPO activity in the samples was estimated by interpolation from the standard curve.

#### Elastase activity

Elastase activity was determined by the chromogenic substrate N-Methoxysuccinyl-Ala-Ala-Pro-Val p-nitroanilide (M4765, Sigma-Aldrich, USA). Supernatants were used for measuring elastase activity. Briefly, 20 µl of supernatant was added to 80 µl of 1 mM substrate in a 96-well transparent, flat bottom plate, and absorbance was measured at 405 nm for 60 minutes. Standard samples with different elastase concentrations (from human leukocytes, E8140, Sigma-Aldrich, USA) were prepared to obtain a standard curve in the range of 16-166 mU/ml. Elastase activity in the samples was estimated by interpolation from the standard curve.

Assuming that only neutrophils produce myeloperoxidase and elastase enzymes, the enzyme content per neutrophils was calculated using the following formula:

96 well reaction volume = V = 150 µl, Total number of cells in volume (V) = A, Percentage of neutrophils in a sample (from flow cytometry) = X, Total number of Neutrophils in volume (V) = A.X, Total enzyme in reaction volume = B (Interpolated from the standard curve). Total enzyme per neutrophil = B/(A.X)

### Mathematical model

A mechanistic mathematical model was developed based on existing knowledge of granulopoiesis(13) and the following assumptions. Assumptions: (i) Differentiation of granulocyte-monocyte progenitors (GMPs) to myelocytes occurs through several intermediate stages (such as myeloblast and promyelocyte), with each stage having a specific proliferation rate. The model assumes that the proliferating pool is kinetically homogeneous with identical turnover rates for all the stages. (ii) The numbers of senescent neutrophils returning to bone marrow again for clearance(16) are assumed to be negligible. (iii) Both in non-inflammatory and the induced inflammatory condition, it is assumed that a steady-state is established, and the labeled cells are tracked at the steady-state. (iv) As the half-life of EDU in circulation (around half an hour)(17) is much lower than the timescale of the experiment, mitotic pool neutrophil progenitors/precursors are assumed to be labeled instantaneously. Equations describing the percentage of EDU positive neutrophils in various compartments:

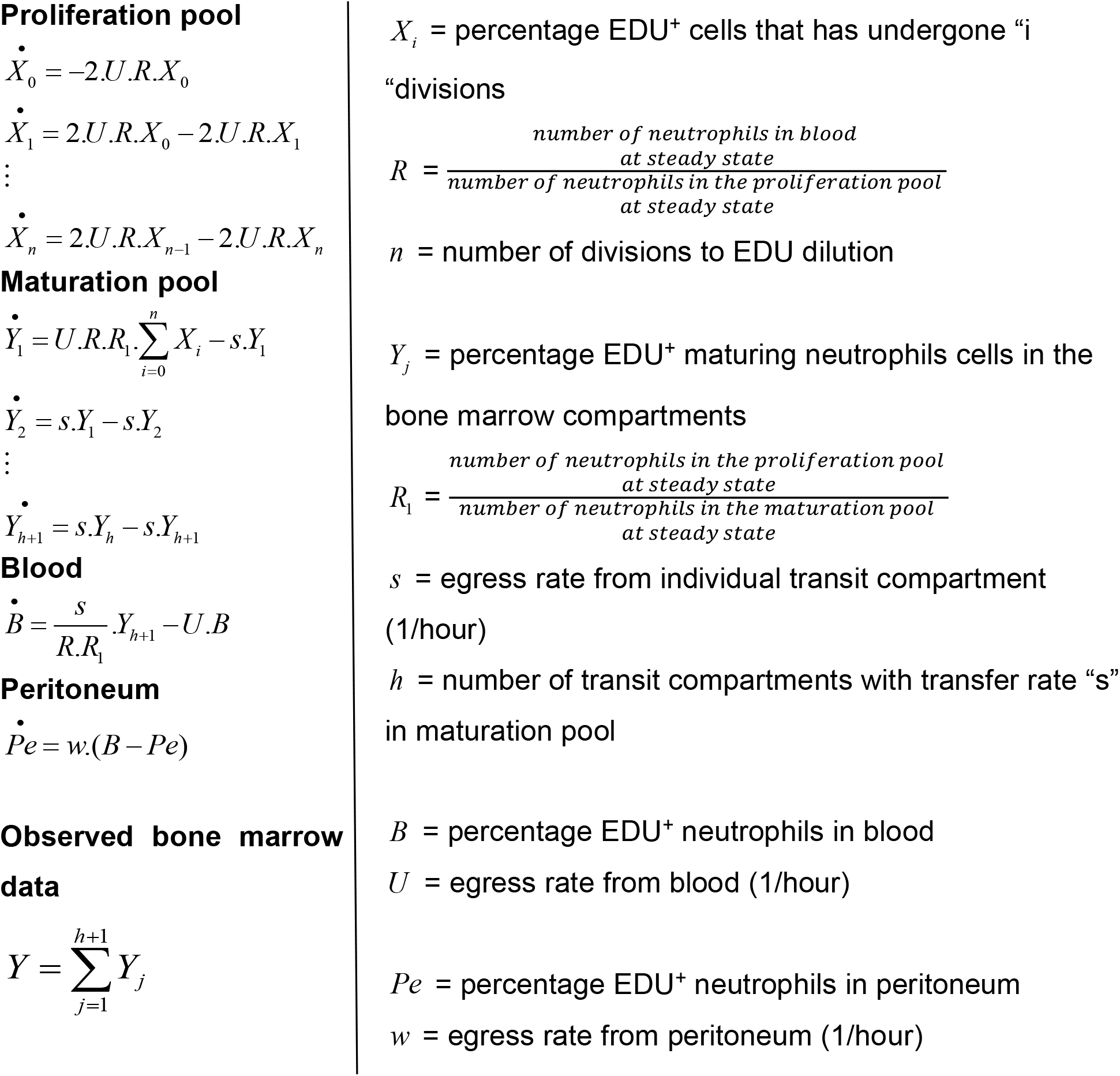

The detailed derivation of the equations is provided in the Supplementary Information.

### Parameter estimation

Agreement between model estimation and experimental data is quantified(18) as given below, where noise in the experimental measurement is assumed to be normally distributed:

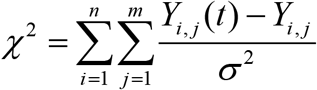

n = number of datasets

m = number of time points

Y_i,j_ = measured data

Y_i,j_(t) = estimated value

Equations are numerically solved using ODE45 in MATLAB with initial conditions that all compartments except proliferation pool has no EdU tagged cells initially (at t=0). The initial percentage of EDU tagged cells, **α** in the proliferation pool, is a parameter in the model. The distribution of parameters yielding acceptable agreement between model and data are obtained using iterative Approximate Bayesian Computation (iABC)(19) and iABC procedure was implemented as described(20). SI and observability analysis were performed analytically (see supplementary information).

### Statistics

All data presented are based on at least 2 or more independent experiments with at least 3 animals per experimental group. An independent experiment is described as an experiment involving new/different batches of microspheres, mice and performed on a different date. Each ‘n’ represents an individual animal. Data were analyzed, and graphs generated using GraphPad Prism 8 (GraphPad Software, La Jolla, CA, USA). One-way ANOVA was used for all statistical comparisons involving multiple groups. For the neutrophil tracking data across different tissue sites (BM, Blood and PC), one-way ANOVA was used at each time point. Significance is represented as * p<0.05, ** p<0.01, *** p<0.001 and **** p<0.0001. Data are presented as mean ± standard deviation.

Common Language Effect Size (CLES) was used to determine statistical differences in parameters estimated using the mathematical model. The CLES measure gives us the probability of one random sample drawn from one distribution has a higher value than a randomly drawn sample from another distribution. CLES measure was calculated using the MATLAB implementation(21) and is reported with 95% confidence interval.

## RESULTS

### Biomaterial implant model for inducing inflammation

Mouse peritoneal cavity (PC) has been used as a model to study the molecular and cellular events for initiation, persistence, and resolution of inflammation(22). Some of the most commonly used irritants (zymosan, LPS and thioglycolate), however, result in an inflammatory microenvironment that lasts for short (<3 days) periods(22–24). We have previously shown that peritoneal implantation of sterile biomaterial microspheres results in sustained inflammation, which may be used to study neutrophil production under inflammatory conditions(25). Hence, mice received sterile biomaterial microspheres or saline as a control (referred to as mock). To tune the level of inflammation, we used chitosan (highly stimulatory(26) in its crude form) and ultrapure alginate (less stimulatory(14, 25)) to prepare the biomaterial microspheres (**Fig. S2**). Implantation of these biomaterial microspheres resulted in recruitment of neutrophils to the implantation site for at least ten days, with a concomitant increase in the percentage of these cells (**Fig. S3**), as reported previously(14, 25).

As a measure of the level of inflammation, we determine the proportions of granulocyte-monocyte progenitors (GMP), which increase when an inflammatory stimulus induces emergency granulopoiesis(27). GMP population was identified as Lineage^-^ c-Kit^+^ Sca-1^-^ CD16/32^+^ CD34^+^ cells through flow cytometry (**Fig. S4**). We observe that in mice implanted with alginate microspheres, the percentages (and number) of granulocyte-monocyte progenitors (GMP) in both the bone marrow and spleen were similar to those in mock controls at seven- and ten-day post-implantation (**Fig. 1A**). In contrast, chitosan microsphere implantation significantly increased GMP percentages in bone marrow and spleen (**Fig. 1A**).

**Figure 1.**
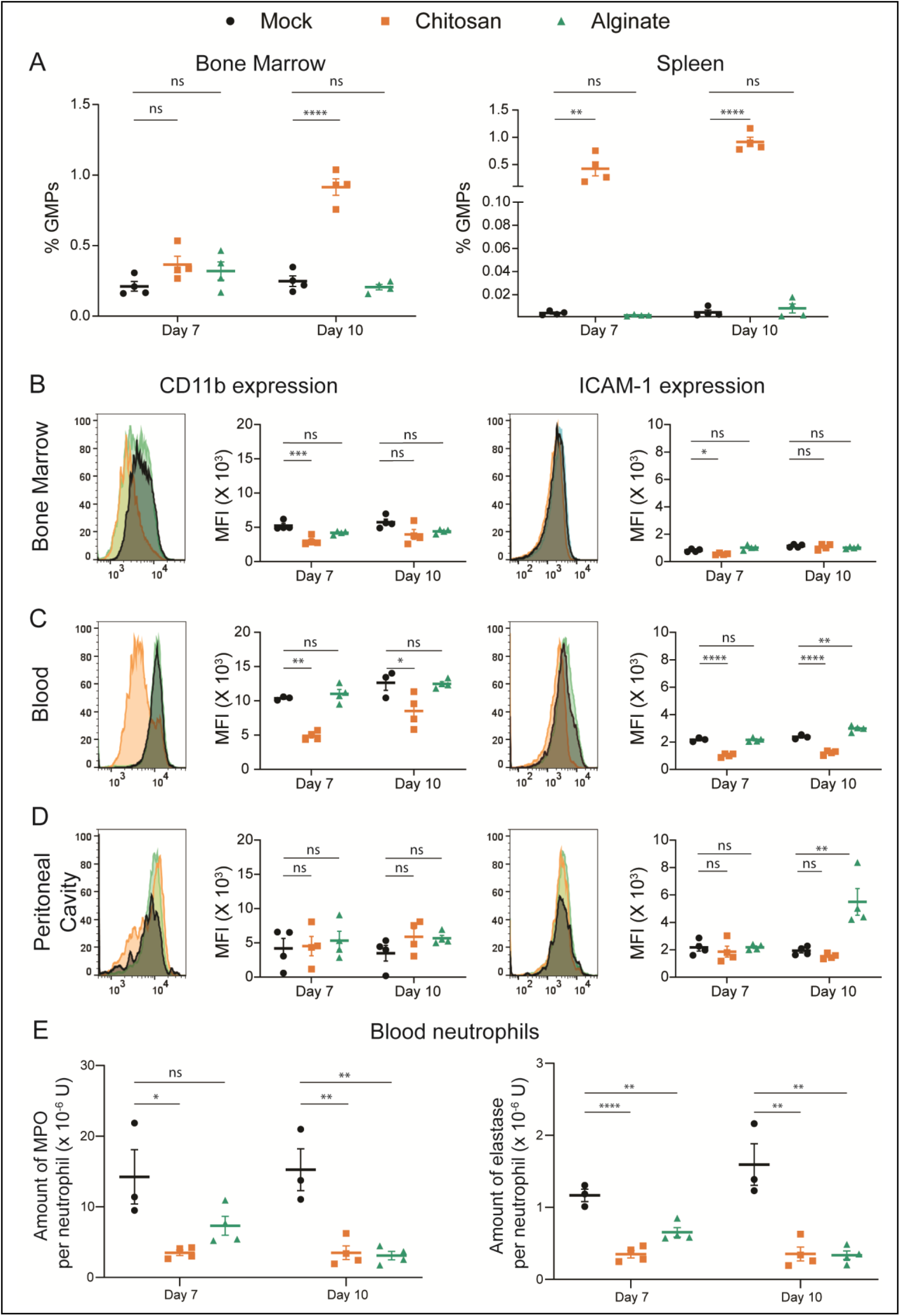
Progenitor numbers and neutrophil phenotype post implantation of sterile biomaterial microspheres. **A** – Quantification of granulocyte-monocyte progenitors (GMPs) as a percentage of total live single cells from bone marrow (BM) and spleen at day 7 and 10 after mock or microsphere-implantation procedure. **B-D** – Phenotype of neutrophils at different tissues, at day 7 and 10 after mock or microsphere-implantation procedure. CD11b and ICAM-1 expression levels among neutrophils following ex vivo activation in BM **(B)**, blood **(C)** and peritoneal cavity **(D). E** – Total intracellular MPO and elastase content per neutrophil in the blood. Saline and chitosan group is representative of 2 independent experiments with total n = 4 mice per group. Alginate group is representative of 1 independent experiment with n = 4 mice at each time point, involving both male and female mice. For statistical analyses, a one-way ANOVA was performed followed by Tukey post-test, and * = p < 0.05; ** = p < 0.01; *** = p < 0.001; and **** = p < 0.0001.

To further understand the effects of microsphere implantation, we studied the phenotype and granular content of neutrophils isolated from bone marrow, blood and PC (implant site). For phenotyping, we measured the surface expression levels of CD11b, ICAM-1, CD62L and CXCR2 on neutrophils at seven- and ten-day post-implantation. The expression levels of these proteins are expected to vary based on the level of cellular activation(28). We observed that in neutrophils retrieved from the bone marrow and blood of mice implanted with chitosan microspheres, expression of CD11b and ICAM-1 following *ex vivo* activation is significantly lower compared to cells from mice implanted with alginate microspheres or mock controls (**Fig. 1B and 1C**). These differences are not observed among neutrophils at the site of inflammation (**Fig. 1D**). Additionally, activation-dependent upregulation of CD11b and ICAM-1 expression was significantly reduced among neutrophils from mice implanted with chitosan microspheres (**Fig. S5**). Activation-dependent downregulation of CD62L and CXCR2 expression was also compromised in neutrophils isolated from the blood of mice implanted with chitosan microspheres as compared to those from mice with alginate microspheres or mock controls (**Fig. S6**). The granular content of neutrophils from the blood of mice implanted with microspheres was also different relative to mock controls. Significantly lower amounts of myeloperoxidase (MPO) and elastase per neutrophil were observed in the blood of mice with microsphere implants (**Fig. 1E**).

Overall, increased GMP percentage in the bone marrow and spleen, and altered surface protein expression and reduced granule enzyme content among neutrophils in circulation suggest that emergency granulopoiesis (EG)-like conditions are induced following chitosan microsphere implantation. However, as GMP levels remains similar in mice with alginate microspheres and mock controls, we label the former as having caused inflammation but not inducing emergency granulopoiesis.

### Neutrophil Kinetics

We next assessed the kinetics of neutrophils under non-inflammatory (mock), inflammatory-(alginate), and inflammation-inducing EG-like (chitosan) conditions. 5-ethynyl-2-deoxyuridine (EdU) labeled neutrophils were identified through the gating strategy described in **Fig. 2A**. Under non-inflammatory and inflammatory (but not EG-like) conditions, labeled neutrophil peaks were observed at day E2 in bone marrow and day E3-E4 in the blood, which is line with recently published data on neutrophils kinetics under steady-state conditions(6). In the presence of inflammation that induces EG-like conditions, we observed that labeled neutrophils appeared earlier in the bone marrow (at day E1, **Fig. 2B**) and blood (at day E2, **Fig. 2C**) as compared to the other two conditions. Percentages of labeled neutrophils in the PC (**Fig. 2D**) followed a trend similar to those in circulation. Under EG-like conditions, the peak of labeled neutrophils appeared at day E2-E3, which was earlier than the peak observed in the other two conditions (day E3-E4). The differences in the timing of the peak of labeled neutrophils at each tissue site in the mice with EG-like conditions suggest a faster release of neutrophils into the circulation and early arrival of these labeled cells at the inflammatory site.

**Figure 2.**
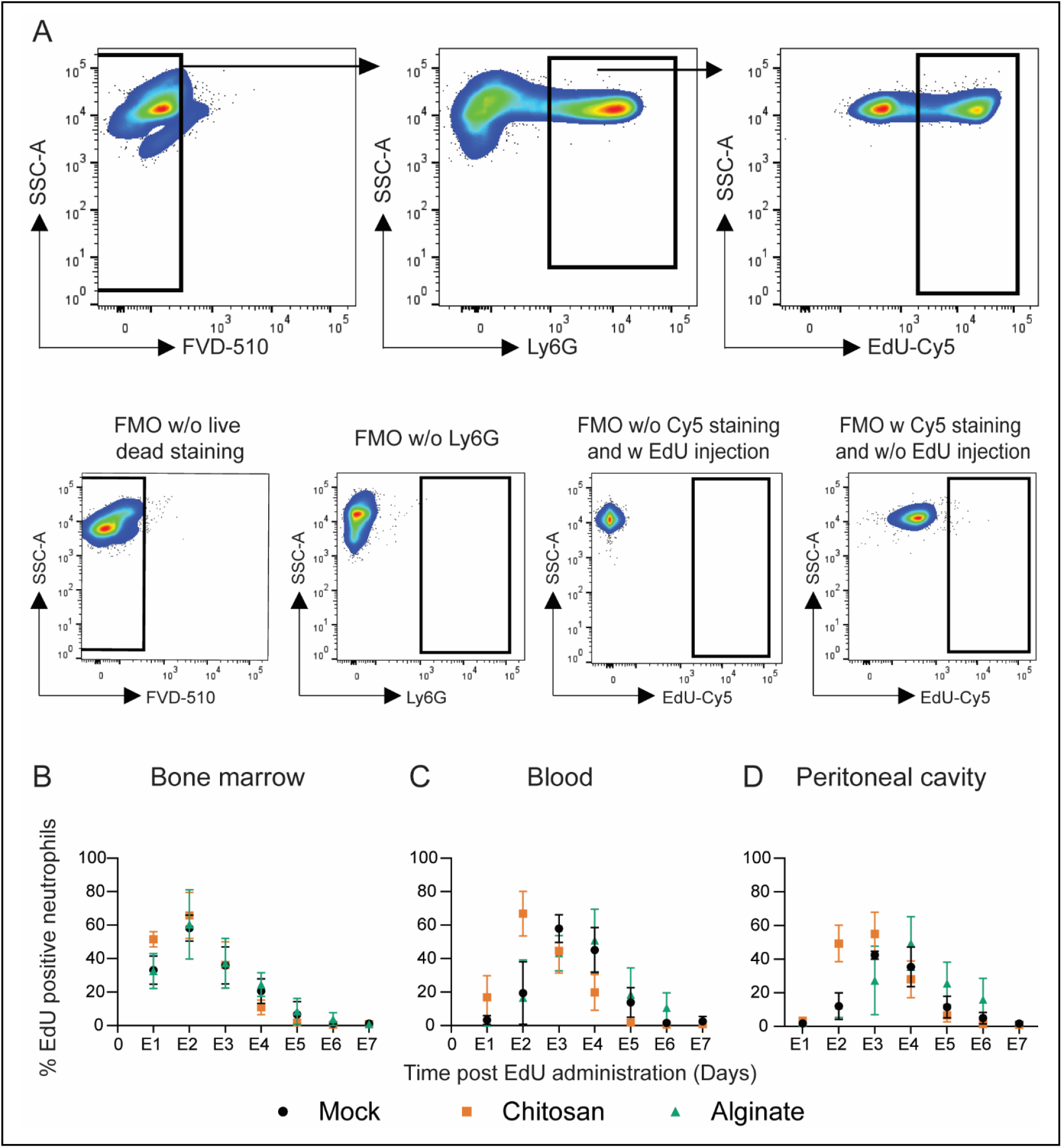
*In vivo* tracking of labeled neutrophils. **A** -Gating strategy used to determine labeled neutrophils. Fluorescence-minus-one (FMO) plots were used to ascertain gate positions. Plots are representative of multiple independent experiments. **B-D** – Labeled neutrophil (EdU positive) percentages in bone marrow **(B)** blood **(C)** and peritoneal cavity **(D)** at different times. N = 3-7 mice/time point/per group pooled from at least 3 independent experiments including both male and female mice. For statistical analyses, at each time point, a one-way ANOVA was performed followed by Tukey post-test. Statistical analysis is summarized in **Table S1**.

Basu et al.(12) previously described differences in neutrophil kinetics based on the extent of stain incorporation. We analyzed our data similarly by separating the EdU incorporated neutrophils as either high expression of EdU (neutrophil incorporating EdU in their last stages of division) or medium expression of EdU (neutrophils that have undergone few divisions after EdU incorporation). Similar to the observations from total EdU-labeled cells, EdU-high and EdU-medium cells appeared earlier in the blood and PC of mice implanted with chitosan microspheres, compared to other conditions (**Fig. S7**). Additionally, we observed a one-day delay in EdU-medium cells’ peaks compared to EdU-high cells in all groups. This delay is expected as the latter are cells arising from progenitors in their last stages of division, leading to a faster appearance in the maturation pool and circulation(12).

The overall time taken for labeled neutrophils to appear and then disappear may also be analyzed to provide insights into residence time in various compartments. One method often employed to understand half-life and residence times is fitting an exponential decay curve from the peak(6, 12). However, an exponential decay function is not an actual depiction of the underlying process and may lead to inaccurate estimation of the parameters. Therefore, we developed a mechanistic model to extract kinetic parameters from the experimental data.

### Quantifying Kinetic Parameters

We applied a system of differential equations that explicitly model EDU dilution, neutrophil maturation in the bone marrow, and residence times in the blood and tissue **(Fig. S8**). The model parameters must be Structurally Identifiable (which is a consequence of the model topology) for them to be estimated(29, 30). By analyzing the transfer functions of the experimentally observed variables and the observability of internal state variables, we determine that the model parameters were identifiable (supplementary information). Then, we fit the model to the data obtained from mock controls and microsphere implanted mice, beginning with uniform priors for all the parameters in iterative Approximate Bayesian Computation (iABC). Representative fits (∼1000) of the model to data (**Fig. 3**) and the overall distribution of each parameter (25000 parameter sets under acceptable chi-square cut-off) are shown (**Fig. 4**).

**Figure 3.**
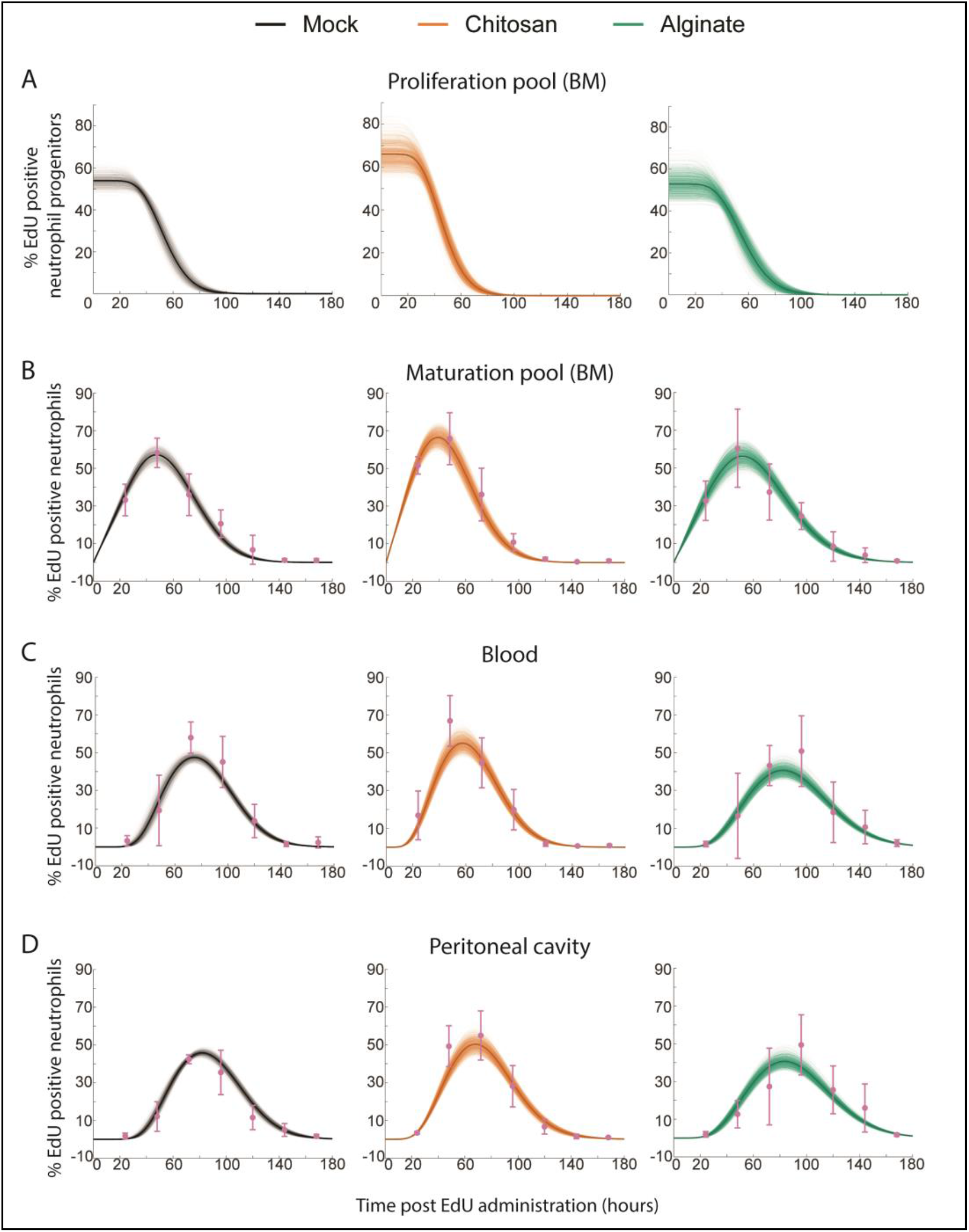
Fitting the mathematical model to experimental data. Representative fits to mock, chitosan and alginate data. Sample curves corresponding to different parameters sets in proliferation pool **(A)**, maturation pool **(B)**, blood **(C)** and peritoneal cavity **(D)** are shown as thin lines and average is shown as thick solid line. Experimentally measured data are shown as pink dots with standard deviatio

**Figure 4.**
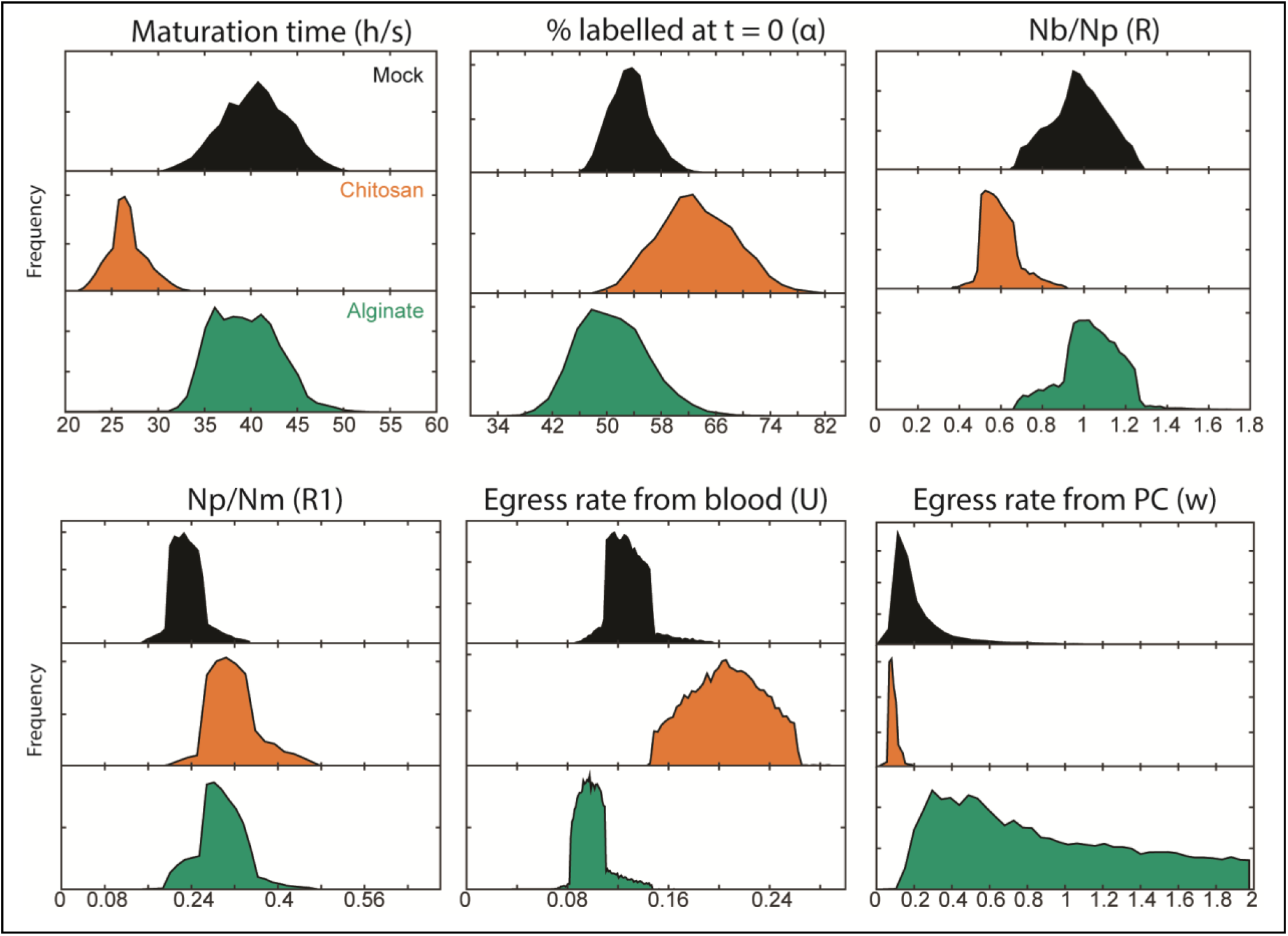
Distributions of parameters obtained from the model. At the end of final iteration of iABC, the possible numerical values for each parameter are provided as a frequency distribution. Np – Number of neutrophils in the proliferation pool at steady state, Nb – Number of neutrophils in the blood at steady state, Nm – Number of neutrophils in the maturation pool at steady state.

The practical identifiability of the parameter (which depends on the discrete nature and quality of the measured data) appears as the magnitude of spread in the parameter distributions(31, 32). This spread is presented in **Table 1** as the medians and 95% confidence intervals of each parameter. Using these parameters, we calculated the values for maturation time in the bone marrow, and half-life and residence time in the blood and PC (**Table 2**). Owing to the large sample size in parameter distributions, comparing the distributions between different groups will show statistical significance even if the differences are very small. To look at practical significance of the differences, which would not be greatly influenced by the large sample size, a non-parametric effect size measure (Common Language Effect Size (CLES)), was used to quantify the differences(33). CLES was calculated between different groups for all the parameters and tabulated (**Table S2**).

**Table 1.** Rate parameters estimated from the mathematical model. Median and 95% confidence interval (ci) of the possible values the parameters can take based on our numerical simulation are tabulated.

|  | Mock |  |  |  | Chitosan |  |  |  | Alginate |  |  |
| --- | --- | --- | --- | --- | --- | --- | --- | --- | --- | --- | --- |
|  | median | ci |  |  | median | ci |  |  | median | ci |  |
| **n**<br>(number of divisions to EDU dilution) | **12** | 10 | 16 |  | **10** | 8 | 14 |  | **11** | 8 | 14 |
| **α**<br>(initial percentage of labelled cells) | **53.37** | 48.16 | 60.13 |  | **62.84** | 52.53 | 75.28 |  | **50.43** | 41.95 | 61.88 |
| **U (hour<sup>-1</sup>)**<br>(egress rate from blood) | **0.12** | 0.10 | 0.16 |  | **0.20** | 0.15 | 0.25 |  | **0.09** | 0.08 | 0.13 |
| **R**<br>(Number of neutrophils in blood / Number of neutrophils in proliferation pool) | **0.97** | 0.70 | 1.23 |  | **0.58** | 0.46 | 0.83 |  | **1.03** | 0.73 | 1.25 |
| **R1**<br>(Number of neutrophils in proliferation pool / Number of neutrophils in maturation pool) | **0.23** | 0.17 | 0.31 |  | **0.31** | 0.240 | 0.43 |  | **0.29** | 0.20 | 0.39 |
| **s (hour<sup>-1</sup>)**<br>(egress rate from individual transit compartment) | **0.29** | 0.23 | 0.34 |  | **0.16** | 0.14 | 0.19 |  | **0.12** | 0.10 | 0.16 |
| **h**<br>(number of transit compartment with transfer rate “s” in maturation pool) | **12** | 9 | 15 |  | **5** | 4 | 5 |  | **5** | 4 | 6 |
| **w (hour<sup>-1</sup>)**<br>(egress rate from peritoneum) | **0.17** | 0.08 | 1.41 |  | **0.08** | 0.06 | 0.16 |  | **0.77** | 0.20 | 1.91 |

**Table 2.** Calculated parameters of neutrophil kinetics. From the rate parameters presented in table 1, parameters of interest were calculated and are presented here as median and 95% confidence interval (ci) of the possible values the parameters can take.

|  | Mock |  |  |  | Chitosan |  |  |  | Alginate |  |  |
| --- | --- | --- | --- | --- | --- | --- | --- | --- | --- | --- | --- |
|  | median | ci |  |  | median | ci |  |  | median | ci |  |
| **Maturation time in bone marrow** (h/s)<br>(hours) | **40.50** | 33.27 | 47.40 |  | **26.50** | 22.87 | 31.34 |  | **39.23** | 33.82 | 46.55 |
| **Half-life in blood**<br>(ln(2)/U)<br>(hours) | **5.46** | 4.089 | 6.81 |  | **3.35** | 2.67 | 4.55 |  | **7.04** | 5.14 | 8.36 |
| **Residence time in blood** (1/U)<br>(hours) | **7.88** | 5.89 | 9.83 |  | **4.84** | 3.85 | 6.57 |  | **10.17** | 7.42 | 12.06 |
| **Half-life in peritoneum**<br>(ln(2)/w)<br>(hours) | **3.99** | 0.49 | 8.10 |  | **8.05** | 4.10 | 11.03 |  | **0.89** | 0.36 | 3.38 |
| **Residence time in peritoneum**<br>(1/w)<br>(hours) | **5.76** | 0.70 | 11.69 |  | **11.61** | 5.91 | 15.92 |  | **1.29** | 0.52 | 4.87 |

Through these steps, we determined that in mice with inflammation that induces EG-like conditions, the neutrophil maturation time in the bone marrow is considerably lower compared to the other two conditions. While this was apparent from the experimental data shown in Fig. 2 (peak of EdU labelled neutrophils in blood appearing at day 2 in mice with chitosan microspheres), the mathematical model allowed us determine that the median of the range of maturation times was lowered to 26 hours (compared to the expected 40 hours under non-inflammatory conditions). Additionally, we observed that half-life (and consequently residence time) in blood decreased, while half-life in PC increased by two-fold in mice with EG-like conditions compared to non-inflammatory conditions. This was, in fact, not apparent from the experimental data.

Together, these data suggest that in response to inflammation resulting in EG-like conditions, neutrophils are generated and released faster from the bone marrow and spend less time in the blood and more time at the site of inflammation when compared to a non-inflammatory condition. In contrast, under a milder inflammatory condition (alginate implants), maturation time is not altered, but the half-life of neutrophils in the blood increases modestly.

## DISCUSSION

Neutrophils play an essential role in pathogen and foreign body clearance(34). However, their aberrant activity may result in pathological conditions such as inflammatory bowel disease(35), rheumatoid arthritis(36), and acute respiratory distress syndrome(37). A potential strategy to enable their beneficial roles while also limiting their damaging effects is the modulation of neutrophil frequencies and life cycle(1, 13, 38). To achieve such modulation, we must first determine neutrophil kinetics under inflammatory conditions.

Granulopoiesis under emergency conditions has been explored by injecting granulocyte colony stimulating factor (G-CSF)(11) or *E. coli*(39). Both G-CSF and E. coli injections resulted in neutrophils appearing faster in circulation (11, 39), and it was suggested (but not demonstrated) that this was due to reduced maturation time in the bone marrow. Another study has determined changes in neutrophil production rates in the bone marrow following a burn injury(40), but other aspects of the kinetics have not been determined. Our data on neutrophil kinetics under emergency granulopoiesis-like conditions induced by chitosan microspheres is in agreement with these reports, and demonstrates that maturation time in the bone marrow is indeed reduced. Alterations in maturation time could have implications on neutrophil phenotype and functions. For example, the lowered capacity to upregulate activation markers (such as CD11b) and lower granule protein content per cell could be a consequence of reduced maturation times in mice implanted with chitosan microspheres. We also determine neutrophils kinetics under conditions that induce inflammation but do not appear to cause emergency granulopoiesis (alginate microsphere implantation). Under such conditions, neutrophil maturation times in the bone marrow remain unchanged.

Further, no previous report describes neutrophil kinetics at the site of inflammation. In this context, a recent study by Ballesteros et al.(6) shows that neutrophil half-life in different tissues under steady-state conditions is similar (except in skin) to their half-life in the blood. It has also been suggested that inflammation may extend neutrophils’ lifespan at the inflammatory tissue site(2, 41, 42). Our data show an increase in half-life (and consequently lifespan) of neutrophils at a sterile inflammatory site if the inflammation induces EG-like conditions.

To estimate maturation times and lifespans of neutrophils in various tissues, we have developed a new model to fit the data. Prior studies have fit an exponential decay function (peak to baseline) to similar experimental data(6, 11, 12), but such modeling ignores the simultaneous ingress of labeled cells during that period resulting in over-estimation of half-lives. Additionally, deriving information on specific aspects of kinetics, such as maturation times in the bone marrow, is not possible using exponential decay based fitting methods. Hence, we modeled the process of granulopoiesis and neutrophil presence in tissues mechanistically, to extract kinetic parameters with an acceptable spread in distributions. Our model provided quantifiable values for most parameters, except for the peritoneal egress rate from mice implanted with alginate microspheres. This could be because of the large variability seen in the experimental data in this group.

One caveat to the data and the model presented here are that we assume that the number of reverse-migrating neutrophils(43) in the bone marrow is negligible compared to the numbers present in the maturation compartment. Another shortcoming is that we do not explicitly include the marginated pool of neutrophils present in the lungs and liver. A logical next step to the current study would be to determine how both reverse migration and marginated pool of neutrophils are altered under inflammatory conditions.

Results presented here showcases the possibility of using biomaterials as a tool to study immune cell dynamics(44, 45). Biomaterials induce differing grades of immune responses based on the chemical nature of the material used and on the physical (shape, size, morphology etc.) characteristics of the implant(46, 47). An example of this variation is demonstrated here with alginate and chitosan-based biomaterial implants. While the chemical composition of these materials is dissimilar and the purity of the base material was different, the exact reasons that these materials induce distinct immune responses remains unclear. Nevertheless, it is noteworthy that the implantation of biomaterials (specifically, chitosan) in the peritoneal cavity results in a dramatic change in neutrophil kinetics, which could have implications in our methodologies to test the compatibility of materials for *in vivo* use.

In conclusion, we show that in the presence of inflammation that induces EG-like conditions, neutrophil maturation time in the bone marrow and half-life in the blood reduces, and residence time at the inflammatory site increases when compared to non-inflammatory and low-grade inflammatory stimuli. The functional impact of such changes remain to be evaluated.

## ACKNOWLEDGEMENT

We thank Virta Wagde and Shruthi KS for assistance with animal work and cell culture. We acknowledge the support of the staff at the central animal facility, IISc.

## REFERENCES

1. Ley, K., H. M. Hoffman, P. Kubes, M. A. Cassatella, A. Zychlinsky, C. C. Hedrick, and S. D. Catz. 2018. Neutrophils: New insights and open questions. Sci. Immunol. 3: eaat4579.

2. Summers, C., S. M. Rankin, A. M. Condliffe, N. Singh, A. M. Peters, and E. R. Chilvers. 2010. Neutrophil kinetics in health and disease. Trends Immunol. 31: 318–324.

3. Borregaard, N. 2010. Neutrophils, from Marrow to Microbes. Immunity 33: 657–670.

4. Bardoel, B. W., E. F. Kenny, G. Sollberger, and A. Zychlinsky. 2014. The Balancing Act of Neutrophils. Cell Host Microbe 15: 526–536.

5. Strydom, N., and S. M. Rankin. 2013. Regulation of Circulating Neutrophil Numbers under Homeostasis and in Disease. J Innate Immun 5: 304–314.

6. Ballesteros, I., A. Rubio-Ponce, M. Genua, E. Lusito, I. Kwok, G. Fernández-Calvo, T. E. Khoyratty, E. van Grinsven, S. González-Hernández, J.Á. Nicolás-Ávila, T. Vicanolo, A. Maccataio, A. Benguría, J. L. Li, J. M. Adrover, A. Aroca-Crevillen, J. A. Quintana, S. Martín-Salamanca, F. Mayo, S. Ascher, G. Barbiera, O. Soehnlein, M. Gunzer, F. Ginhoux, F. Sánchez-Cabo, E. Nistal-Villán, C. Schulz, A. Dopazo, C. Reinhardt, I. A. Udalova, L. G. Ng, R. Ostuni, and A. Hidalgo. 2020. Co-option of Neutrophil Fates by Tissue Environments. Cell 183: 1282-1297.e18.

7. Price, T. H., G. S. Chatta, and D. C. Dale. 1996. Effect of recombinant granulocyte colony-stimulating factor on neutrophil kinetics in normal young and elderly humans. Blood 88: 335–340.

8. Lahoz-Beneytez, J., M. Elemans, Y. Zhang, R. Ahmed, A. Salam, M. Block, C. Niederalt, B. Asquith, and D. Macallan. 2016. Human neutrophil kinetics: modeling of stable isotope labeling data supports short blood neutrophil half-lives. Blood 127: 3431–3438.

9. Ng, L. G., R. Ostuni, and A. Hidalgo. 2019. Heterogeneity of neutrophils. Nat Rev Immunol 19: 255– 265.

10. Pillay, J., I. den Braber, N. Vrisekoop, L. M. Kwast, R. J. de Boer, J. A. M. Borghans, K. Tesselaar, and L. Koenderman. 2010. In vivo labeling with 2H2O reveals a human neutrophil lifespan of 5.4 days. Blood 116: 625–627.

11. Lord, B. I., G. Molineux, Z. Pojda, L. M. Souza, J. J. Mermod, and T. M. Dexter. 1991. Myeloid cell kinetics in mice treated with recombinant interleukin-3, granulocyte colony-stimulating factor (CSF), or granulocyte-macrophage CSF in vivo. Blood 77: 2154–2159.

12. Basu, S., G. Hodgson, M. Katz, and A. R. Dunn. 2002. Evaluation of role of G-CSF in the production, survival, and release of neutrophils from bone marrow into circulation. Blood 100: 854–861.

13. Hidalgo, A., E. R. Chilvers, C. Summers, and L. Koenderman. 2019. The Neutrophil Life Cycle. Trends Immunol. 40: 584–597.

14. Jhunjhunwala, S., S. Aresta-DaSilva, K. Tang, D. Alvarez, M. J. Webber, B. C. Tang, D. M. Lavin, O. Veiseh, J. C. Doloff, S. Bose, A. Vegas, M. Ma, G. Sahay, A. Chiu, A. Bader, E. Langan, S. Siebert, J. Li, D.L. Greiner, P. E. Newburger, U. H. von Andrian, R. Langer, and D. G. Anderson. 2015. Neutrophil Responses to Sterile Implant Materials. PLoS ONE 10: e0137550.

15. Zeng, C., F. Pan, L. A. Jones, M. M. Lim, E. A. Griffin, Y. I. Sheline, M. A. Mintun, D. M. Holtzman, and R. H. Mach. 2010. Evaluation of 5-ethynyl-2′-deoxyuridine staining as a sensitive and reliable method for studying cell proliferation in the adult nervous system. Brain Res. 1319: 21–32.

16. Furze, R. C., and S. M. Rankin. 2008. The role of the bone marrow in neutrophil clearance under homeostatic conditions in the mouse. FASEB J. 22: 3111–3119.

17. Cheraghali, A. M., R. Kumar, E. E. Knaus, and L. I. Wiebe. 1995. Pharmacokinetics and bioavailability of 5-ethyl-2’-deoxyuridine and its novel (5R,6R)-5-bromo-6-ethoxy-5,6-dihydro prodrugs in mice. Drug Metab Dispos 23: 223–226.

18. 2007. Numerical recipes: the art of scientific computing, 3rd ed. (W. H. Press, ed). Cambridge University Press, Cambridge, UK?; New York.

19. Beaumont, M. A., W. Zhang, and D. J. Balding. 2002. Approximate Bayesian Computation in Population Genetics. Genetics 162: 2025–2035.

20. Chhajer, H., V. A. Rizvi, and R. Roy. 2020. Life cycle process dependencies of positive-sense RNA viruses suggest strategies for inhibiting productive cellular infection,. Microbiology.

21. Hentschke, H. hhentschke/measures-of-effect-size-toolbox. https://github.com/hhentschke/measures-of-effect-size-toolbox. Accessed February 1, 2021..

22. Ito, Y., H. Kinashi, T. Katsuno, Y. Suzuki, and M. Mizuno. 2017. Peritonitis-induced peritoneal injury models for research in peritoneal dialysis review of infectious and non-infectious models. Ren Replace Ther 3: 16.

23. Rao, T. S., J. L. Currie, A. F. Shaffer, and P. C. Isakson. 1994. In vivo characterization of zymosan-induced mouse peritoneal inflammation. J Pharmacol Exp Ther 269: 917–925.

24. Miyazaki, S., F. Ishikawa, T. Fujikawa, S. Nagata, and K. Yamaguchi. 2004. Intraperitoneal Injection of Lipopolysaccharide Induces Dynamic Migration of Gr-1high Polymorphonuclear Neutrophils in the Murine Abdominal Cavity. Clin. Diagn. Lab. Immunol. 11: 452–457.

25. Jhunjhunwala, S., D. Alvarez, S. Aresta-DaSilva, K. Tang, B. C. Tang, D. L. Greiner, P. E. Newburger, U. H. von Andrian, R. Langer, and D. G. Anderson. 2016. Splenic progenitors aid in maintaining high neutrophil numbers at sites of sterile chronic inflammation. Journal of Leukocyte Biology 100: 253–260.

26. Hoemann, C. D., and D. Fong. 2017. Immunological responses to chitosan for biomedical applications. In Chitosan Based Biomaterials Volume 1 >Elsevier. 45–79.

27. Manz, M. G., and S. Boettcher. 2014. Emergency granulopoiesis. Nat Rev Immunol 14: 302–314.

28. Fortunati, E., K. M. Kazemier, J. C. Grutters, L. Koenderman, and van J. M. M. Van den Bosch. 2009. Human neutrophils switch to an activated phenotype after homing to the lung irrespective of inflammatory disease. Clin. Exp. Immunol. 155: 559–566.

29. Bellman, R., and K.J. Åström. 1970. On structural identifiability. Math. Biosci. 7: 329–339.

30. Villaverde, A. F. 2019. Observability and Structural Identifiability of Nonlinear Biological Systems. Complexity 2019: 1–12.

31. Raue, A., C. Kreutz, T. Maiwald, J. Bachmann, M. Schilling, U. Klingmüller, and J. Timmer. 2009. Structural and practical identifiability analysis of partially observed dynamical models by exploiting the profile likelihood. Bioinformatics 25: 1923–1929.

32. Lehmann, E. L., and G. Casella. 2001. Theory of Point Estimation {Springer Texts in Statistics},. Springer-Verlag New York Inc., Dordrecht.

33. McGraw, K. O., and S. P. Wong. 1992. A common language effect size statistic. Psychol. Bull. 111: 361–365.

34. Mayadas, T. N., X. Cullere, and C. A. Lowell. 2014. The Multifaceted Functions of Neutrophils. Annu. Rev. Pathol. Mech. Dis. 9: 181–218.

35. Zhou, G. X., and Z. J. Liu. 2017. Potential roles of neutrophils in regulating intestinal mucosal inflammation of inflammatory bowel disease: Role of neutrophils in IBD. J. Dig. Dis. 18: 495–503.

36. Wright, H. L., R. J. Moots, and S. W. Edwards. 2014. The multifactorial role of neutrophils in rheumatoid arthritis. Nat. Rev. Rheumatol. 10: 593–601.

37. Yang, S.-C., Y.-F. Tsai, Y.-L. Pan, and T.-L. Hwang. 2020. Understanding the role of neutrophils in acute respiratory distress syndrome. Biomed. J. S2319417020301499.

38. Németh, T., M. Sperandio, and A. Mócsai. 2020. Neutrophils as emerging therapeutic targets. Nat. Rev. Drug Discov. 19: 253–275.

39. Xie, X., Q. Shi, P. Wu, X. Zhang, H. Kambara, J. Su, H. Yu, S.-Y. Park, R. Guo, Q. Ren, S. Zhang, Y. Xu, L. E. Silberstein, T. Cheng, F. Ma, C. Li, and H. R. Luo. 2020. Single-cell transcriptome profiling reveals neutrophil heterogeneity in homeostasis and infection. Nature Immunology 21: 1119–1133.

40. Rosinski, M., M. L. Yarmush, and F. Berthiaume. 2004. Quantitative Dynamics of in Vivo Bone Marrow Neutrophil Production and Egress in Response to Injury and Infection. Ann. Biomed. Eng. 32: 1109–1120.

41. McCracken, J. M., and L.-A. H. Allen. 2014. Regulation of Human Neutrophil Apoptosis and Lifespan in Health and Disease. J. Cell Death 7: 15–23.

42. Filep, J. G., and A. Ariel. 2020. Neutrophil heterogeneity and fate in inflamed tissues: implications for the resolution of inflammation. Am. J. Physiol. Cell Physiol. 319: C510–C532.

43. Nourshargh, S., S. A. Renshaw, and B. A. Imhof. 2016. Reverse Migration of Neutrophils: Where, When, How, and Why? Trends Immunol. 37: 273–286.

44. Jhunjhunwala, S. 2018. Biomaterials for Engineering Immune Responses. J. Indian Inst. Sci. 98: 49– 68.

45. Mariani, E., G. Lisignoli, R.M. Borzì, and L. Pulsatelli. 2019. Biomaterials: Foreign Bodies or Tuners for the Immune Response? Int. J. Mol. Sci. 20.

46. Sadtler, K., A. Singh, M. T. Wolf, X. Wang, D. M. Pardoll, and J. H. Elisseeff. 2016. Design, clinical translation and immunological response of biomaterials in regenerative medicine. Nat Rev Mater 1: 16040.

47. Oakes, R. S., E. Froimchuk, and C. M. Jewell. 2019. Engineering Biomaterials to Direct Innate Immunity. Adv. Therap. 2: 1800157.

