## Supplementary for "Sterile Inflammation Alters Neutrophil Kinetics in Mice"

**This PDF file includes:**

| | |
|---|---|
| Supplementary Text | Pages 2 – 6 |
| Figs. S1 to S8 | Pages 7 – 15 |
| Tables S1 and S2 | Pages 16 – 18 |

**Supplementary Text**

**Model Derivation**

We first write the equations for absolute number of labelled cells in various compartments and later derive equations for percentage labelled cells in each compartment.

Proliferation pool:

The cells labelled in the proliferation pool divide leading to the dilution of EDU content per cell, which eventually results in the label becoming undetectable. EDU dilution in proliferation pool is represented by following equations adopted from Mohri(1).

$$\frac{dx_o}{dt} = -px_o - qx_o$$
$$\frac{dx_1}{dt} = 2px_o - px_1 - qx_1$$
$$\vdots$$
$$\frac{dx_n}{dt} = 2px_{n-1} - px_n - qx_n$$

$x_i$ = number of $EDU^+$ cells that has undergone "i" divisions
$p$ = proliferation rate of homogeneous progenitors (1/hour)
$q$ = egress rate from proliferation pool (1/hour)
$n$ = number of divisions to EDU becoming undetectable

Maturation pool:

One of the possibilities to model maturation delay in bone marrow is using delay differential equations (DDE)(2, 3), but using DDEs would lead to unphysical jumps of labelled neutrophils in blood after a specified delay (Δ), which is not what we observe in the acquired EDU data. Thus, using transit compartments(4) we mechanistically model the maturation to describe the data better. These transit compartments can be thought of as mimicking the actual maturation stages of the neutrophils in maturation pool.

$$\frac{dy_1}{dt} = q\sum_{i=0}^{n} x_i - s \times y_1$$
$$\frac{dy_2}{dt} = s \times y_1 - s \times y_2$$
$$\vdots$$
$$\frac{dy_h}{dt} = s \times y_{h-1} - s \times y_h$$

$y_j$ = number of $EDU^+$ maturing neutrophils cells in the bone marrow compartments
s = egress rate from individual transit compartment (1/hour)
$h$ = number of transit compartments with transfer rate "s" in maturation pool

Mean transit time can be calculated as residence time per transit compartment × number of transit compartments = (1/s) × h.

Blood and Peritoneum:

$$\frac{dz}{dt} = sy_h - Uz$$
$$\frac{dg}{dt} = vz - wg$$

$z$ = number of EDU$^+$ neutrophils in blood
$g$ = number of EDU$^+$ neutrophils in peritoneum
$U$ = egress rate from blood (1/hour)
$v$ = ingress rate into peritoneum (1/hour)
$w$ = egress rate from peritoneum (1/hour)

Steady state constraints and derivation of percentage equations.

Constraints are as follows,

$$p.N_{pr} = q.N_{pr} = s.y_1^{st} = s.y_2^{st} = \cdots = s.y_h^{st} = U.N_b$$

$$v.N_b = w.N_{pe}$$

$$y_1^{st} + y_2^{st} + ... + y_h^{st} = N_{ma}$$

where,
$N_{pr}$ = total number of cells in proliferation pool at steady state
$y_i^{st}$ = total number of neutrophils in i$^{th}$ transit compartment at steady state
$N_b$ = total number of neutrophils in blood at steady state
$N_{pe}$ = total number of neutrophils in peritoneum at steady state
$N_{ma}$ = total number of neutrophils in maturation pool at steady state

Number of cells in each compartment is assumed constant in steady state. Thus, dividing each compartment equations by respective constant steady state numbers will yield the percentage equations as shown in the main text.

**Structural Identifiability**

For linear time invariant systems, such as the one presented here, the SI theory has been completely developed by Bellman and Åström (5). Since the presented model is an open loop system, we were able to derive a general impulse-response transfer function for arbitrary number of compartments, allowing the analysis of the compartment numbers (n, h) as free parameters.

The impulse-response function in Laplace domain of the observed variables are

$$\mathcal{L}\left(Y(t)\right) = Y(S) = \alpha \sum_{j=1}^{h+1} \left[ \frac{s^{j-1}}{\left(S+s\right)^{j}} C \sum_{i=0}^{n} \frac{A^{i}}{\left(S+A\right)^{i}} \right]$$

$$H_{Y,\alpha} = \frac{Y(S)}{\alpha} = \sum_{j=1}^{h+1} \left[ \frac{s^{j-1}}{\left(S+s\right)^{j}} C \sum_{i=0}^{n} \frac{A^{i}}{\left(S+A\right)^{i}} \right]$$

$$H_{B,\alpha} = \frac{\mathcal{L}\left(B(t)\right)}{\alpha} = \frac{U s^{h+1}}{\left(S+s\right)^{h+1}\left(S+U\right)} \sum_{i=0}^{n} \frac{A^{i}}{\left(S+A\right)^{i}}$$

$$H_{Pe,\alpha} = \frac{\mathcal{L}\left(Pe(t)\right)}{\alpha} = \frac{wU s^{h+1}}{\left(S+w\right)\left(S+s\right)^{h+1}\left(S+U\right)} \sum_{i=0}^{n} \frac{A^{i}}{\left(S+A\right)^{i}}$$

where,

$A = 2UR$

$C = URR_1$

For clarity, the transfer functions can be expanded to a fraction with polynomials in numerator and denominator.

$$H_{Y,\alpha} = \frac{C(nh+n+h+1) + \left(\frac{C(Ah^2(n+1)+n^2 s(h+1))}{As}\right)S + \left(\frac{C(A^2 h(n+1)(h-1)^2 + 2Asn^2h^2 + s^2 h(h+1)(h-1)^2)}{2(As)^2}\right)S^2 + \cdots + \left(\frac{C}{A^n s^h}\right)S^{h+n}}{As + \left(\frac{(A+s(n+1))}{As}\right)S + \left(\frac{A^2 h(h+1) + As(n+1)(h+1) + s^2 n(n+1)}{(As)^2}\right)S^2 + \cdots + \left(\frac{1}{A^{n+1}s^{h+1}}\right)S^{h+n+2}}$$

$$H_{B,\alpha} = \frac{U(n+1) + \left(\frac{Un^2}{A}\right)S + \left(\frac{Un(n-1)^2}{2A^2}\right)S^2 + \cdots + \left(\frac{U}{A^n}\right)S^n}{UA + \left(\frac{UA+Us(n+1)+As}{s}\right)S + \left(\frac{A^2(s+Uh^2+Uh) + s^2(n+1)(A+Un) + AsU(n+1)(h+1)}{As^2}\right)S^2 + \cdots + \left(\frac{1}{A^n s^{h+1}}\right)S^{h+n+3}}$$

$$H_{Pe,\alpha} = \frac{Uw(n+1) + \left(\frac{Uwn^2}{A}\right)S + \left(\frac{Uwn(n-1)^2}{2A^2}\right)S^2 + \cdots + \left(\frac{Uw}{A^n}\right)S^n}{UAw + \left(\frac{UwA+Uws(n+1)+UAs}{s}\right)S + \left(\frac{UA^2(s(w+1)+(h^2+h)) + Us^2(n+1)(w(A+Un)+A) + As(wU^2(n+1)(h+1)+As)}{As^2}\right)S^2 + \cdots + \left(\frac{1}{A^n s^{h+1}}\right)S^{h+n+4}}$$

The coefficients of the numerator and denominator polynomials are the structural invariants (6) of the transfer functions and are uniquely determined (7). This implies that the coefficients $f(\overline{P})$ are independent nonlinear equations in $\overline{P}$, where $\overline{P}$ denotes the model parameters.

Non identifiability stems from number of parameters being larger than the available nonlinear constraints (underdetermined system) and consequently leading to infinite solutions. In our case we end up with an overdetermined system of nonlinear equations. The possibility of no solutions is discarded since by design of SI analysis we ask if synthetically generated continuous data by one set of parameters can be reproduced by other sets. Therefore, there is at least one set of parameters determining structural invariants and possible number solutions must be finite in an overdetermined system. Thus, we conclude that the model is identifiable.

**Observability analysis**

Using observability analysis of the proposed model, we show that the inferred initial condition is also identifiable. $EDU^+$ cells in proliferation pool are the internal state variable which are not experimentally observed, but, using observability analysis, we can check if it is possible to infer the internal state variable given an output variable (observed variable). Consider the system in which all the compartments in the bone marrow including both proliferation and maturation transit compartments are state variables and the observed $EDU^+$ cells in maturation pool is the output variable.

Generally, the state variables and the observables can be represented as

$$\frac{d\vec{X}}{dt} = K\vec{X}$$

$$\vec{Y} = M\vec{X}$$

Where $\vec{X}$ is the state variable and $\vec{Y}$ is the observable variable (output).

For the given set of equation for bone marrow (proliferation pool and maturation pool compartments), we have

$$
K=\begin{array}{c}
\begin{array}{cccccc|ccccc}
1 & 2 & 3 & \cdots & n & n+1 & 1 & 2 & 3 & \cdots & h+1
\end{array}\\
\left[\begin{array}{cccccc|ccccc}
-A & 0 & 0 & \cdots & 0 & 0 & 0 & 0 & 0 & \cdots & 0\\
A & -A & & & & 0 & 0 & 0 & 0 & \cdots & 0\\
0 & A & -A & & & 0 & 0 & 0 & 0 & \cdots & 0\\
 & & \ddots & \ddots & & & & & & \vdots & \\
0 & 0 & 0 & \ddots & -A & 0 & 0 & 0 & 0 & & 0\\
0 & 0 & 0 & \cdots & A & -A & 0 & 0 & 0 & \cdots & 0\\
\hline
C & C & C & \cdots & C & C & -s & 0 & 0 & \cdots & 0\\
0 & & \cdots & & 0 & 0 & s & -s & 0 & \cdots & 0\\
0 & & \cdots & & 0 & 0 & 0 & s & -s & \cdots & 0\\
 & & \vdots & & & & & & \ddots & \ddots & \\
0 & & \cdots & & 0 & 0 & 0 & 0 & 0 & s & -s
\end{array}\right]
\begin{array}{c}
1\\ 2\\ 3\\ \vdots\\ n\\ n+1\\ 1\\ 2\\ 3\\ \vdots\\ h+1
\end{array}
\end{array}
$$

$$
M=\begin{array}{c}
\begin{array}{cccccccccccc}
1 & 2 & 3 & \cdots & n & n+1 & 1 & 2 & 3 & \cdots & h
\end{array}\\
\left[\begin{array}{cccccccccccc}
0 & 0 & 0 & \cdots & 0 & 0 & 1 & 1 & 1 & \cdots & 1
\end{array}\right]
\end{array}
$$

According to the observability condition (8),

$$
\mathrm{O}^{L}=\begin{pmatrix} M \\ M.K \\ M.K^{2} \\ \vdots \\ M.K^{n+h+1} \end{pmatrix}
$$

the observability matrix Should have full rank, for given parameters in K and M, for the state variables to be observable.

In the model presented here the state variables are structurally observable (observable for arbitrary parameter realization) since there exists a structurally equivalent model (9, 10) This observability condition means that there is only one curve for $X_n$ that corresponds to output Y(t). Analytical solution for $X_n$ can be written as

$$X_n=\alpha\times\frac{(2pt)^n}{n!}\times e^{-2pt}$$

, where $\alpha$ is the initial percentage of labelled cells.

It follows from the above equation that $\alpha$ is uniquely inferred from output since $X_n$ corresponds uniquely to Y(t). In other words, by fitting the output variable to data, we can infer the percentage of EDU tagged cells in proliferation pool at time t=0.

### Supplementary Figures

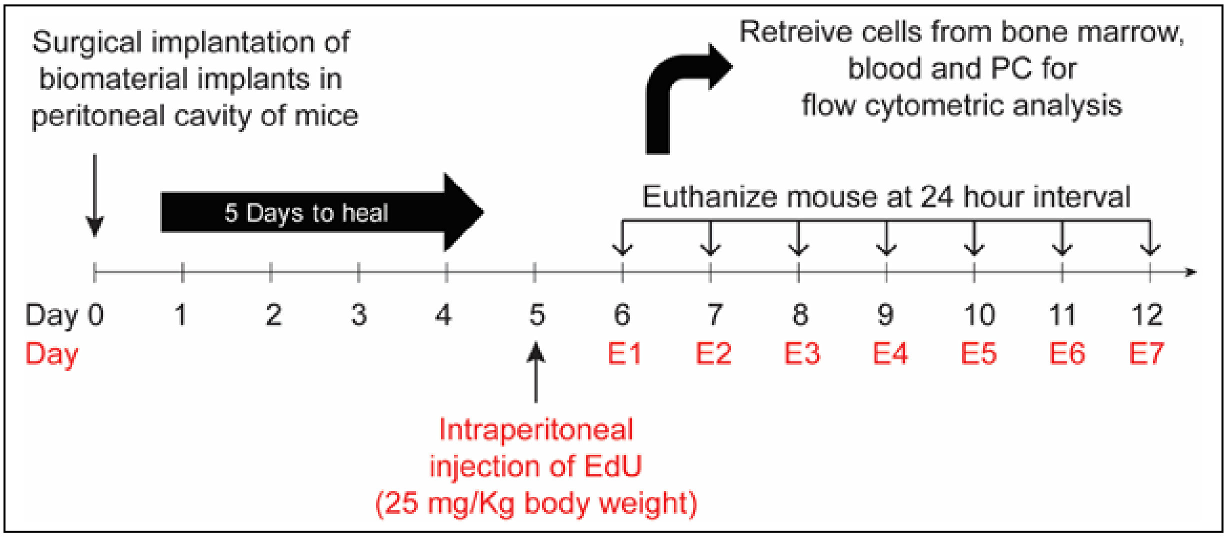


**Fig. S1. Experimental design**. Microspheres were implanted in the peritoneal cavity of mice. At day 5, EdU (5-ethynyl-2-deoxyuridine) was injected intraperitoneally to label proliferating cells. Following EdU administration, at 24-hour intervals, mice were euthanized, and labelled neutrophils were quantified using flow cytometry.

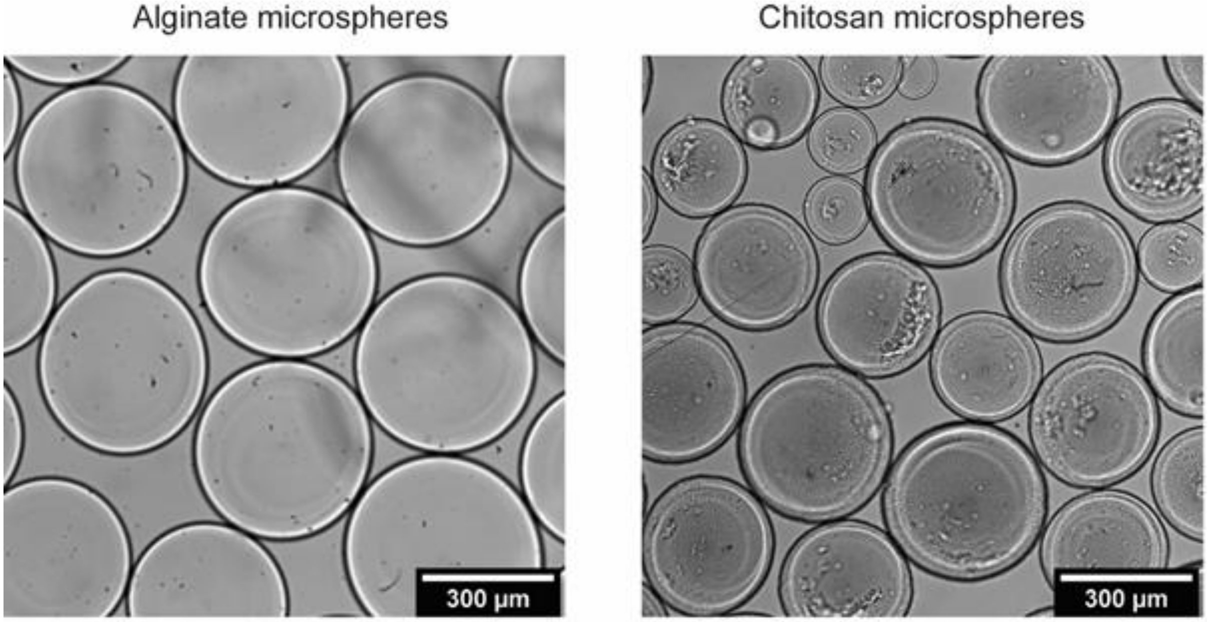


| Biomaterial | Size | Number of microspheres implanted per mouse |
|---|---|---|
| Alginate | 353 ± 41 microns | 1900 ± 690 |
| Chitosan | 337 ± 47 microns | 2185 ± 975 |

**Fig. S2. Sterile implants**. **Top** – Bright field images of alginate and chitosan microspheres. **Bottom table** – Sizes and counts of microspheres implanted in mice presented as mean ± standard deviation. Mice were implanted with ~450 µl of microspheres in all experiments. Total number of microspheres was calculated assuming a packing efficiency of 0.7

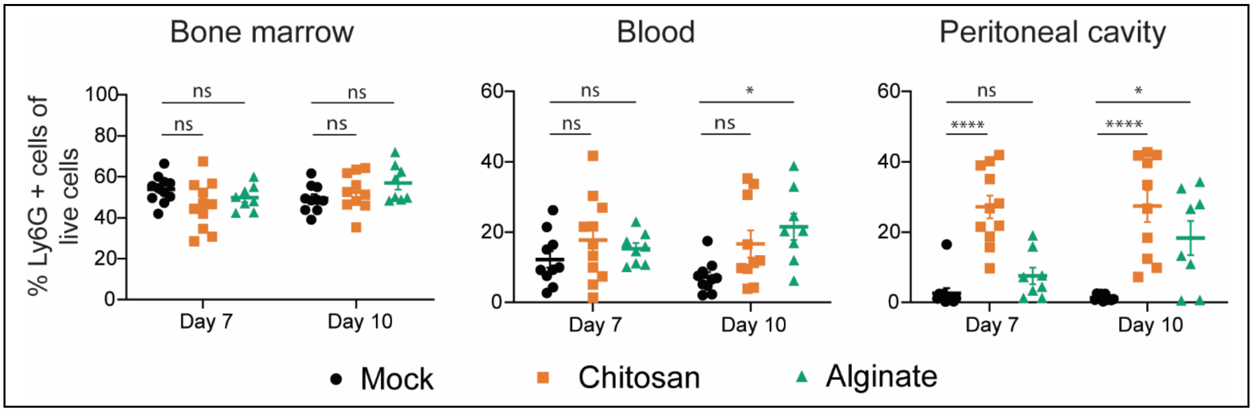


**Fig. S3. Neutrophil frequencies**. Percentage of neutrophils in the bone marrow (BM), blood and peritoneal cavity (PC) after seven and ten days of microspheres implantation. n = 8-11 mice/time point/per group pooled from at least 5 independent experiments. Both male and female mice are included in this study. For statistical analyses, at each time point, a one-way ANOVA was performed followed by Tukey post-test, where * = $p < 0.05$; ** = $p < 0.01$; *** = $p < 0.001$; and **** = $p < 0.0001$.

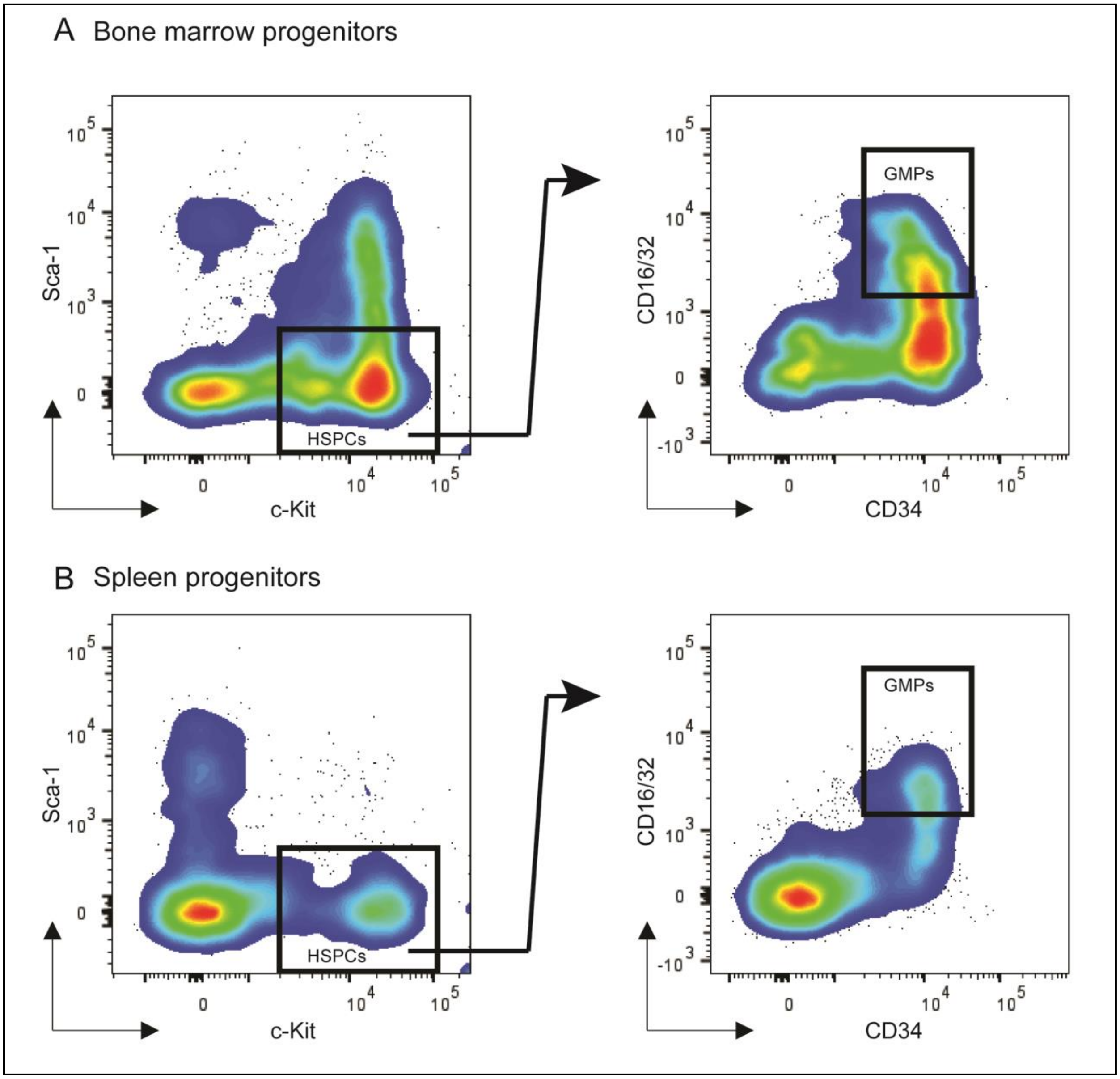


**Fig. S4. Identification of granulocyte-monocyte progenitor (GMP) population**. Representative flow cytometry dot plots to determine GMPs (c-Kit$^{+}$ Sca1$^{-}$ CD34$^{+}$ CD16/32$^{+}$) in the bone marrow **(A)** and spleen **(B).** c-Kit$^{+}$ Sca-1$^{-}$ population was identified as a sub-population of live cells and lineage negative (Lin$^{-}$) cells, where Lin$^{-}$ indicates negative for I-A/I-E, CD49b, NK-1.1, CD11c, Ly6C, Ly6G, CD3, CD127, CD19 and CD45R/B220.

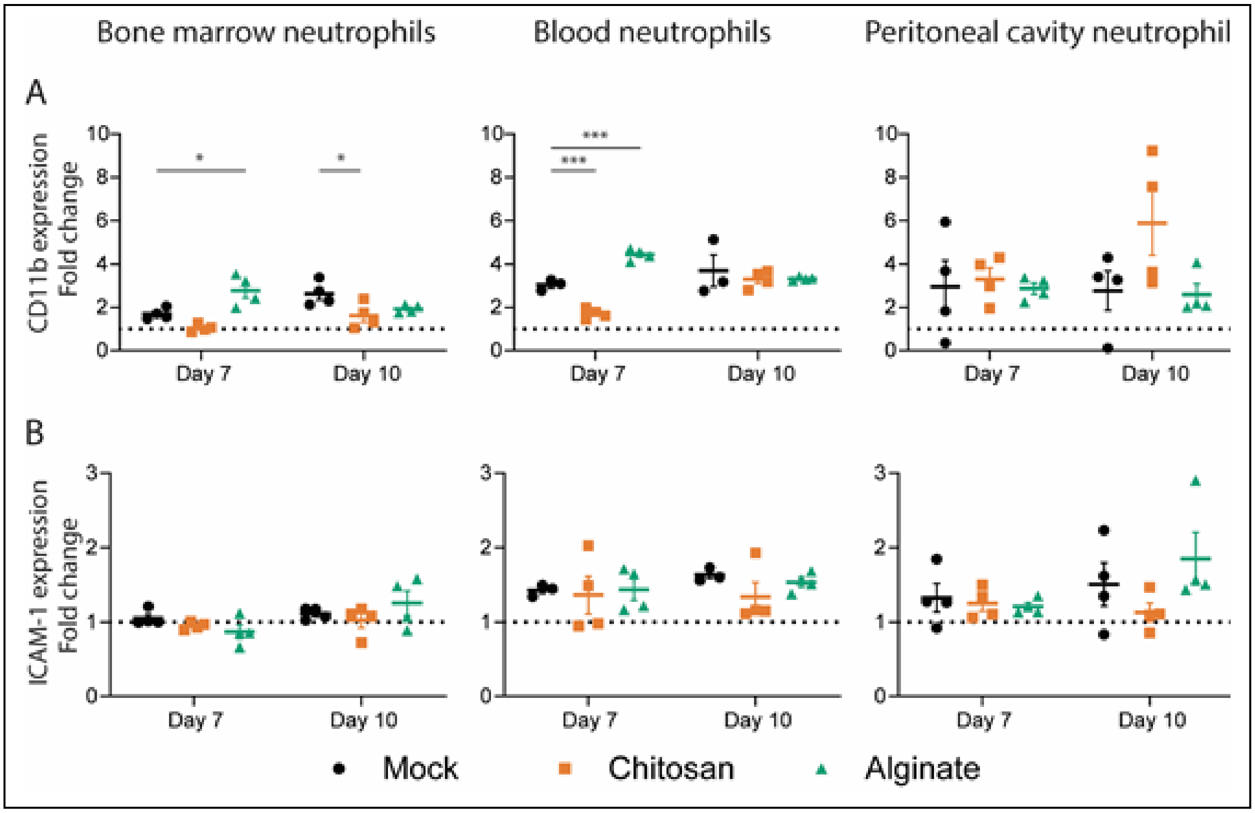


**Fig. S5. Upregulation of CD11b and ICAM-1 upon *ex-vivo* activation.** Cells isolated from the bone marrow (BM), blood and peritoneal cavity (PC) were either treated (activation) or not treated (non-activated) with cytochalasin B (5 µg/ml) and fMLP (5 µM) ex-vivo, and expression levels of CD11b and ICAM-1 among neutrophils ($Ly6G^+$ cells) was quantified. Fold change of CD11b **(A)** and ICAM-1 **(B)** is calculated by dividing MFI of activated neutrophils by non-activated neutrophils. Saline and chitosan groups are representative of 2 independent experiments with total n = 4. Alginate group is representative of 1 independent experiment with n = 4 mice at each time point, involving both male and female mice. For statistical analyses, a one-way ANOVA was performed followed by Tukey post-test, and * = $p < 0.05$; ** = $p < 0.01$; *** = $p < 0.001$; and **** = $p < 0.0001$. Black – Mock, Orange – Chitosan, Green - Alginate

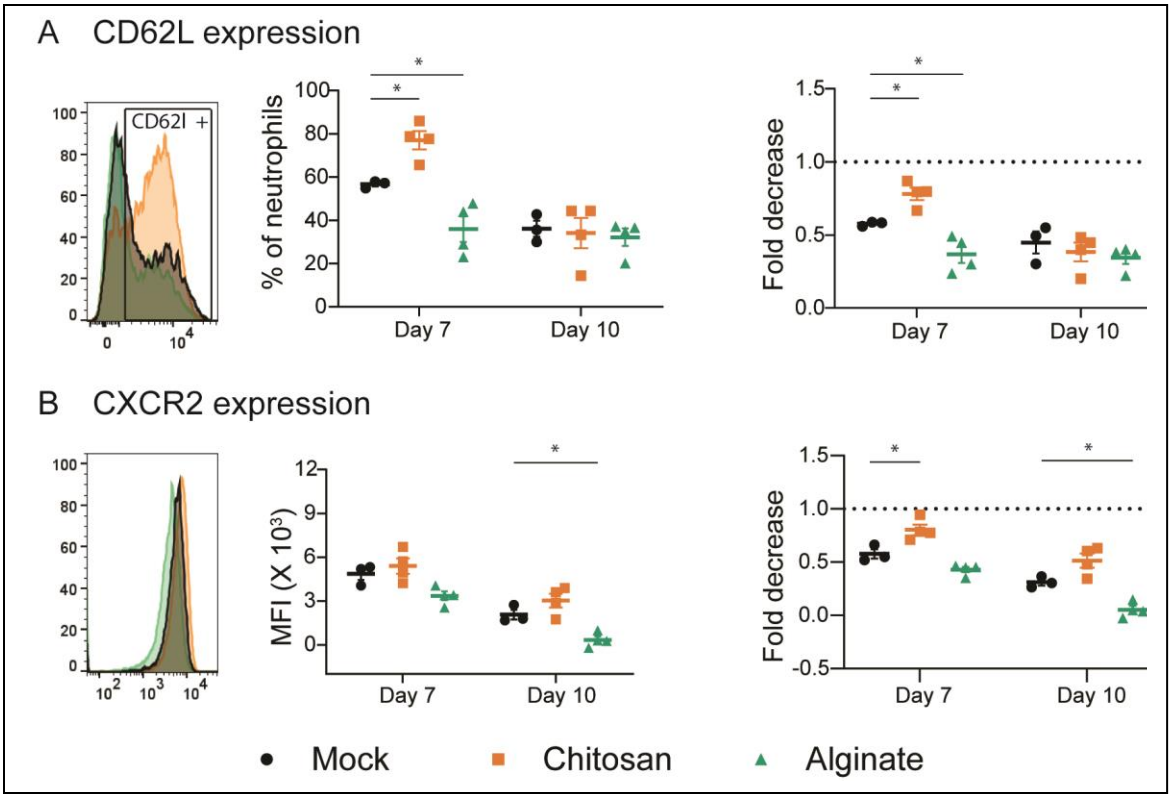


**Fig. S6. CD62L and CXCR2 expression on neutrophils in circulation.** Expression levels of CD62L **(A)** and CXCR2 **(B)** on neutrophils (Ly6G$^+$ cells) was quantified at day 7 and 10 after mock or microsphere-implantation procedure. Left panel is a representative histogram of expression. Center panel is quantification of % of neutrophils expressing CD62L or MFI of CXCR2 following activation. Right panel is fold change in expression following activation. The activation procedure involving addition of cytochalasin B (5 µg/ml) and fMLP ( 5 µM). Fold change was calculated by dividing percentage or MFI post-activation by values under non-activated conditions. Saline and chitosan groups are representative of 2 independent experiments with total n = 4. Alginate group is representative of 1 independent experiment with n = 4 at each time point, involving both male and female mice. For statistical analyses, a one-way ANOVA was performed followed by Tukey post-test, and * = $p < 0.05$; ** = $p < 0.01$; *** = $p < 0.001$; and **** = $p < 0.0001$. Black – Mock, Orange – Chitosan, Green - Alginate

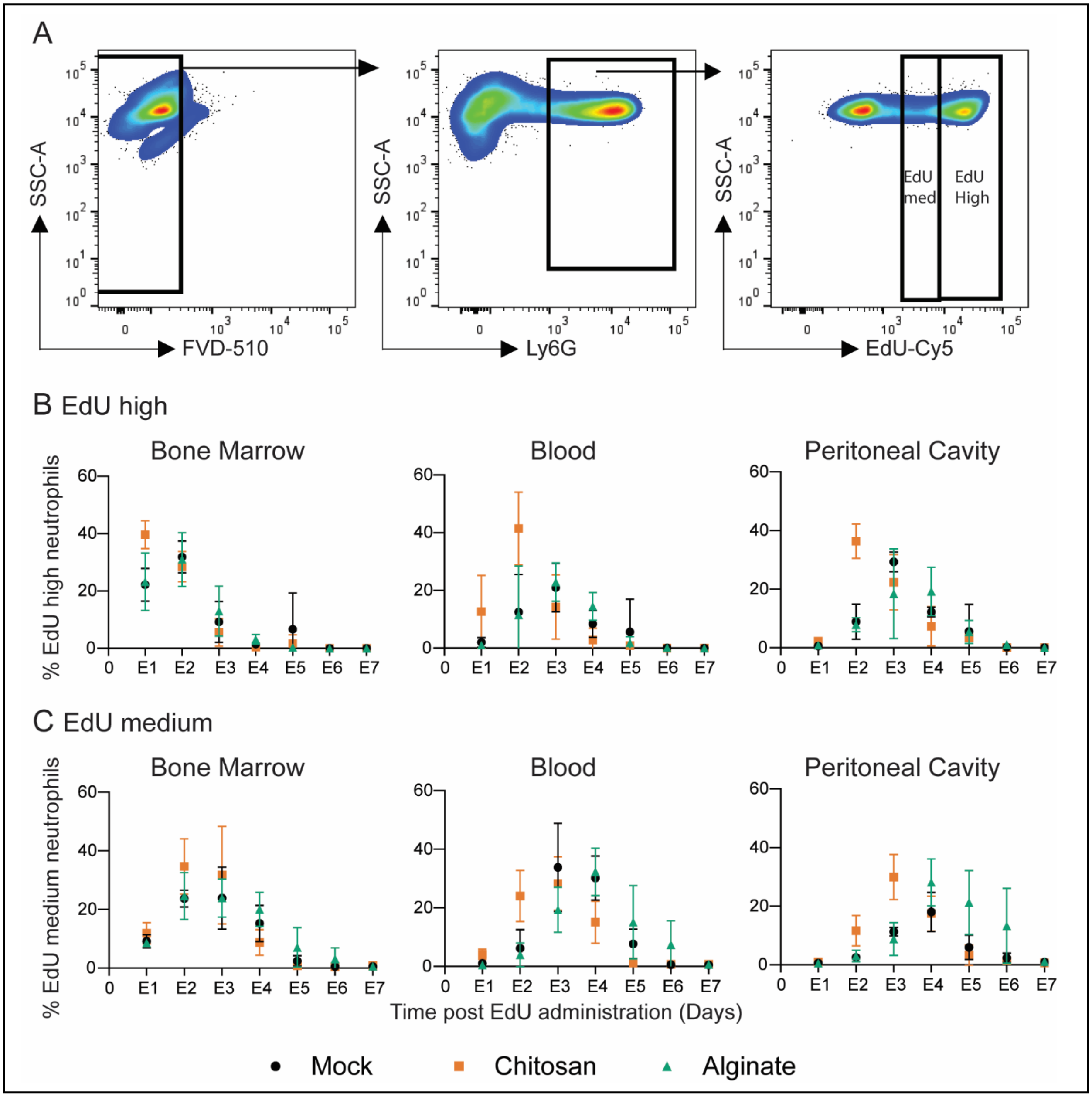


**Fig. S7. *In vivo* tracking of EdU-high and EdU-medium neutrophils. A** - Gating strategy used to quantify EdU-high and EdU-medium neutrophils. Plots are representative of multiple independent experiments. EdU high **(B)** and EdU medium **(C)** neutrophils in the bone Marrow, blood and peritoneal cavity. N = 3-7 mice/time point/per group pooled from at least 3 independent experiments including both male and female mice

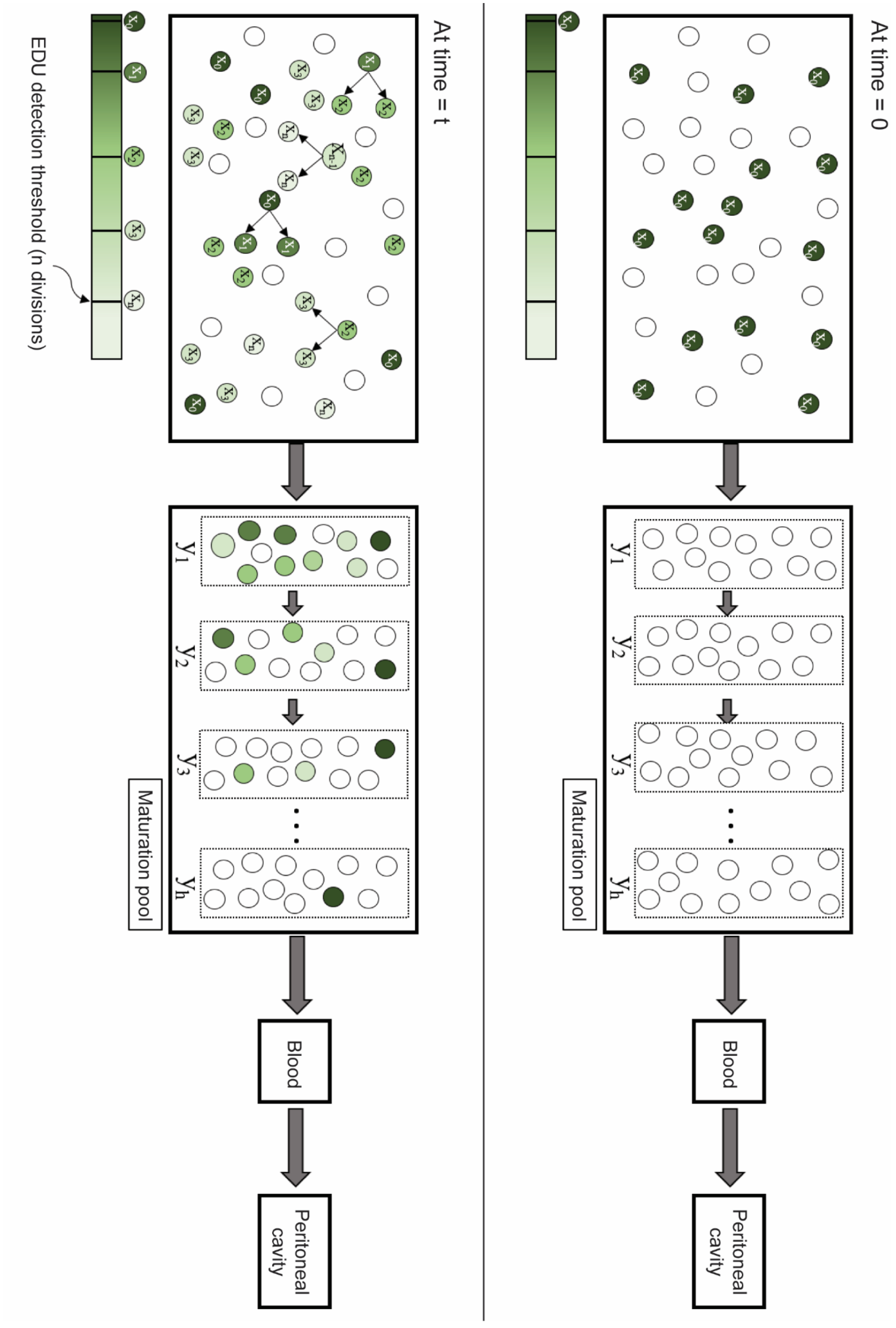
At time = 0
At time = t
Maturation pool
Blood
Peritoneal cavity
EDU detection threshold (n divisions)

**Fig. S8. Schematic of EdU incorporation in the bone marrow.** A depiction of our model showing how EdU labeled neutrophils change over time, in the mitotic pool and the maturation pool, in the bone marrow. The schematic shows $EdU^+$ neutrophils at time t = 0 and at arbitrary time ‘t’.

**Table S1. Statistical analysis of data presented in figure 2.** One-way ANOVA followed by Tukey post-test, and * = $p < 0.05$; ** = $p < 0.01$; *** = $p < 0.001$; and **** = $p < 0.0001$

| | Mock vs Chitosan | | | Mock vs Alginate | | | Alginate vs Chitosan | | |
|---|---|---|---|---|---|---|---|---|---|
| | BM | Blood | PC | BM | Blood | PC | BM | Blood | PC |
| Day E1 | * | * | ns | ns | ns | ns | * | * | ns |
| Day E2 | ns | *** | **** | ns | ns | ns | ns | ** | **** |
| Day E3 | ns | ns | ns | ns | ns | ns | ns | ns | * |
| Day E4 | * | * | ns | ns | ns | ns | ** | ** | * |
| Day E5 | ns | ns | ns | ns | ns | * | ns | * | ** |
| Day E6 | ns | ns | ns | ns | ns | ns | ns | ns | ns |
| Day E7 | ns | ns | ns | ns | ns | ns | ns | ns | ns |

**Table S2. Effect size measures between parameter distributions of different groups.**

| | Mock vs Chitosan | | | | Mock vs Alginate | | | | Alginate vs Chitosan | | |
|---|---|---|---|---|---|---|---|---|---|---|---|
| | CLES | ci | | | CLES | ci | | | CLES | ci | |
| **n** no. of divisions to EDU dilution | 0.200 | 0.198 | 0.206 | | 0.240 | 0.236 | 0.244 | | 0.444 | 0.439 | 0.449 |
| **α** initial percentage of labelled cells | 0.929 | 0.927 | 0.931 | | 0.329 | 0.324 | 0.334 | | 0.943 | 0.941 | 0.945 |
| **U** (hour$^{-1}$) egress rate from blood | 0.988 | 0.985 | 0.989 | | 0.083 | 0.077 | 0.093 | | 0.99 | 0.99 | 0.99 |
| **R** no. of neutrophils in blood / no. of neutrophils in proliferation pool | 0.0108 | 0.0101 | 0.0114 | | 0.6146 | 0.6097 | 0.6194 | | 0.0043 | 0.0040 | 0.0046 |
| $\mathbf{R_1}$ no. of neutrophils in proliferation pool /no. of neutrophils in maturation pool | 0.9362 | 0.9344 | 0.9382 | | 0.8759 | 0.8727 | 0.8789 | | 0.6443 | 0.6396 | 0.6488 |
| **s** (hour$^{-1}$) egress rate from individual transit compartment | 6E-5 | 5.25E-5 | 6.85E-5 | | 1.23E-7 | 9.68E-8 | 1.55E-7 | | 0.965 | 0.963 | 0.966 |
| **h** no. of transit compartment with transfer rate “s” in maturation pool | 2.17E-7 | 1.75E-7 | 2.67E-7 | | 3.26E-6 | 2.76E-6 | 3.89E-6 | | 0.255 | 0.251 | 0.259 |
| **w** (hour$^{-1}$) egress rate from peritoneum | 0.267 | 0.265 | 0.269 | | 0.840 | 0.836 | 0.843 | | 0.059 | 5.8E-2 | 6.09E-2 |
| **s/h** (hours) maturation time | 4.94E-4 | 4.49E-4 | 5.39E-4 | | 0.421 | 0.416 | 0.426 | | 9.1E-4 | 8.47E-4 | 9.80E-4 |

### REFERENCES


1. Mohri, H. 1998. Rapid Turnover of T Lymphocytes in SIV-Infected Rhesus Macaques. *Science* 279: 1223–1227.

2. Pillay, J., I. den Braber, N. Vrisekoop, L. M. Kwast, R. J. de Boer, J. A. M. Borghans, K. Tesselaar, and L. Koenderman. 2010. In vivo labeling with 2H2O reveals a human neutrophil lifespan of 5.4 days. *Blood* 116: 625–627.

3. Lahoz-Beneytez, J., M. Elemans, Y. Zhang, R. Ahmed, A. Salam, M. Block, C. Niederalt, B. Asquith, and D. Macallan. 2016. Human neutrophil kinetics: modeling of stable isotope labeling data supports short blood neutrophil half-lives. *Blood* 127: 3431–3438.

4. Kahn, M. G., L. M. Fagan, and L. B. Sheiner. 1991. Combining physiologic models and symbolic methods to interpret time-varying patient data. *Methods Inf Med* 30: 167–178.

5. Bellman, R., and K. J. Åström. 1970. On structural identifiability. *Mathematical Biosciences* 7: 329–339.

6. Vajda, S. 1981. Structural equivalence of linear systems and compartmental models. *Mathematical Biosciences* 55: 39–64.

7. Boas, R. P. 1935. A Proof of the Fundamental Theorem of Algebra. *The American Mathematical Monthly* 42: 501.

8. R. Kalman. 1959. On the general theory of control systems. *IRE Transactions on Automatic Control* 4: 110–110.

9. Barnett, S. 1978. Dynamical Systems and their Applications. Linear Theory. By J. L. CASTI. (New York: Academic Press, 1977) [Pp. 240] Price£13.85. *International Journal of Control* 27: 983–983.

10. DiStefano, J. J. 2013. *Dynamic systems biology modeling and simulation*, First edition. Elsevier, Academic Press, Amsterdam.
